# Functional Genomic Screens Reveal RBBP4 as a Key Regulator of Cell Cycle Progression in TMZ-Resistant Glioblastoma

**DOI:** 10.1101/2025.05.09.653010

**Authors:** Ezgi Yagmur Kala, Filiz Senbabaoglu Aksu, Erdem Ercan, Altar Ozbiyik, Ahmet Cingöz, Ozlem Yedier-Bayram, Ali Cenk Aksu, Ayse Derya Cavga, Ebru Yilmaz, Ipek Kok, A. Humeyra Dur Karasayar, Ibrahim Kulac, Hamzah Syed, Martin Philpott, Adam P. Cribbs, Tugba Bagci-Onder

## Abstract

Temozolomide (TMZ) remains the standard of care for glioblastoma; however, its efficacy is frequently influenced by epigenetic mechanisms, notably the methylation status of the O6-methylguanine-DNA methyltransferase (MGMT) promoter. While MGMT promoter hypermethylation is associated with enhanced responsiveness to TMZ, additional epigenetic determinants of TMZ resistance remain largely undefined. In this study, we established TMZ-resistant glioblastoma cell lines that consistently maintained their resistant phenotype both *in vitro* and *in vivo*. Transcriptomic analyses revealed a marked upregulation of MGMT expression in these models. To systematically investigate the epigenetic regulators governing TMZ resistance and cell survival, we conducted CRISPR/Cas9-based functional genomic screens using our focused Epigenetic Knock-Out Library (EPIKOL), which targets 800 chromatin regulators alongside selected positive and negative controls. These unbiased screens validated MGMT as a primary mediator of TMZ resistance, confirming the robustness of our approach. Moreover, dropout screens across multiple resistant cell line models identified Retinoblastoma Binding Protein 4 (RBBP4) as a critical vulnerability. Notably, RBBP4 knockout significantly impaired cell proliferation without affecting MGMT expression, suggesting a distinct mechanism supporting the survival of TMZ-resistant glioblastoma cells. Subsequent transcriptomic profiling following RBBP4 loss demonstrated significant downregulation of cell cycle pathways, particularly the G2/M checkpoint. Live-cell imaging and immunofluorescence analyses further revealed increased cell size and multinucleation in RBBP4-deficient cells, indicative of disrupted mitotic progression. Collectively, our results identify RBBP4 as a key regulator of cell cycle progression and survival in TMZ-resistant glioblastoma and highlight its potential as a novel epigenetic target for therapeutic intervention in recurrent disease.

## INTRODUCTION

Glioblastoma is the most common and lethal primary brain tumor, classified as Grade 4 by the World Health Organization (WHO)(1). The prognosis for glioblastoma remains poor, with a 5-year survival rate of 7.1%, and the incidence rate increases depending on age and sex (2). Standard treatment involves surgical resection, radiation, and chemotherapy, where Temozolomide (TMZ) is the first-line chemotherapeutic agent for treatment but unfortunately recurrence is very common (3,4).

TMZ is an oral, small, lipophilic DNA alkylating agent used as the primary treatment for glioblastoma (3). Once inside the cell, TMZ is hydrolyzed into 5-(3-methyltriazen-1-yl) imidazole-4-carboxamide (MTIC), which adds methyl groups to DNA bases N^3^-methyladenine, N^7^-methylguanine, and O^6^-methylguanine (5). Cells with active DNA repair mechanisms such as Mismatch Repair (MMR) and Base Excision Repair (BER), or with high O^6^-methylguanine methyltransferase (MGMT) enzyme activity, can repair this damage, leading to TMZ resistance. In contrast, cells with defective repair pathways undergo apoptosis (6–9). As MGMT enzyme is responsible for the repair of O^6^-methylguanine lesions, its promoter methylation status serves as a key prognostic factor in glioblastoma (10,11). Patients with hypermethylated *MGMT* promoters exhibit reduced MGMT expression, leading to improved TMZ response and longer survival (11). Conversely, unmethylated *MGMT* results in TMZ resistance due to higher MGMT activity and repair of TMZ-induced DNA damage (12,13). Efforts to overcome TMZ resistance include MGMT inhibitors, such as O^6^-benzylguanine, however significant side effects underscore the need for further research to identify more effective treatments (14,15).

Epigenetic modifications are frequently observed in glioblastoma, contributing to tumor progression and therapeutic resistance (16). With the updated WHO classifications, glioblastomas are recognized as Isocitrate dehydrogenase (*IDH*)-wild type tumors (1), which are now distinguished at epigenetic level from *IDH*-mutant gliomas exhibiting a hypermethylated DNA phenotype (17). Along with the accumulating knowledge on the epigenetic alterations as drivers of glioblastoma progression, chromatin modifiers are increasingly recognized as therapeutic targets. For example, increased expression of several classes of histone deacetylases (HDACs) are associated with poor prognosis, such as HDAC4 and HDAC6 from Class II HDACs, which have been associated with radioresistance in glioblastoma (18,19). Therefore, HDAC inhibitors have been prime candidates in clinical trials, mostly in combination with chemotherapeutic agents or radiation therapy, such as Vorinostat (SAHA) (16). Similarly, histone demethylases (KDMs) are recognized as clinically important for glioblastoma and *IDH*-mutant glioma (20). Intriguingly, KDM5A has been shown to regulate TMZ response (21,22). Overall, epigenetic modifications play a crucial role in glioblastoma progression and resistance, emphasizing the unmet need for uncovering novel therapeutic strategies to overcome resistance and treat recurrent tumors. To this end, interrogating the function of all chromatin modifiers, such as the writer, eraser and reader complexes is a powerful approach to isolate specific epigenetic intervention points in glioblastoma. Using functional genomics screening approach with our epigenetic focused CRISPR/Cas9 libraries, we previously identified novel epigenetic regulators of tumor cell fitness in several cancers including glioblastoma (23,24). However, the epigenetic adaptations at the therapy-resistant tumor cell state remained elusive.

In this study, we generated multiple *in vitro* models of acquired TMZ resistance to investigate the epigenetic factors underlying chemotherapy resistance. Functional experiments confirmed the resistance phenotype both *in vitro* and *in vivo*, and transcriptomics analysis revealed that TMZ-resistance was accompanied by activated MGMT expression. Applying functional EPIKOL CRISPR screens to examine epigenetic vulnerabilities of TMZ-resistant cells, we identified Retinoblastoma Binding Protein 4 (*RBBP4*) as a master regulator of TMZ-resistant cell fitness. RBBP4, also known as RbAp48, is a histone chaperone identified in association with many chromatin regulating complexes (25). In contrast with the previous findings that implicate *RBBP4* in DNA repair mechanisms in therapy-naïve glioblastoma cells (26–28). Our results provide novel insights into the pleiotropic functions of epigenetic regulators in therapy-resistant models, which can open up new therapeutic options for recurrent glioblastoma.

## METHODS

### Cell culture

Human glioblastoma cell lines U87MG, T98G and U373 along with Human Embryonic Kidney 293T cells, were purchased from American Type Culture Collection (ATCC). Normal Human Astrocytes (NHA) were purchased from ScienCell (USA). All cell lines were cultured in DMEM (Gibco, USA), supplemented with 1% Penicillin/Streptomycin (Biowest, USA) and 10% Fetal Bovine Serum (Biowest, USA). Astrocytes were seeded over culture dishes coated with 0.01 w/v Poly-l-lysine (Sigma-Aldrich, USA). Cells were maintained in humidifier incubators with 5% CO^2^ and 37°C. All cell lines were routinely screened for mycoplasma infection.

### Generation of Temozolomide-resistant glioblastoma cell lines

U87MG cells were treated with Temozolomide (TMZ) (Selleckchem, S1237, USA) with a dose escalation approach, starting with 5 µM. Dose was doubled every 2-3 weeks or when cells show resistance, until 200 µM. U373 cells were subjected to a high dose selection (250 µM) for 15 days with TMZ renewed every day, and the remaining population was expanded. Generated TMZ-resistant cells were named U87MG-TR and U373-TR, respectively. Parental U87MG and U373 cells were treated with DMSO and passaged alongside the resistant cells. All cells were maintained with TMZ (200 µM for U87MG-TR and 250 µM for U373-TR) during culturing.

### Cell viability assay

Cells were seeded as 1000 cells/well in 96-well plates (Sarstedt, Germany). For IC_50_ value determination, cells were treated with TMZ with 1:2 serial dilution (starting from 1000 µM). At day 5, 3 mg/ml MTT reagent (Cayman, China) was added into the wells (1:4 of the initial medium) and plates were incubated for 1-4 hours at 37°C. Medium was discarded and DMSO was added to dissolve the formazan crystals. Absorbance based measurement (570 nm) was performed using Synergy H1 Reader (Biotech, USA). For growth curve assays, cells were seeded at post-transduction day 9 (PT9) and MTT measurements were taken at Day1 (PT10), Day3 (PT12) and Day5 (PT14).

### Colony formation assay

Cells were seeded to 6-well plates (Sarstedt, Germany) as 750-800 cells/well and 350-400 cells/well to 12-well plates (Sarstedt, Germany) as triplicates. TMZ treatment was performed the next day and medium was replaced after 72 hours. For CRISPR/Cas9 assays, cells were seeded for colony formation assay at PT9. Cells were kept in culture for 10-14 days until visible colonies were observed. Afterwards, medium was discarded, cells were fixed with cold 100% Methanol for 5 minutes, followed by Crystal Violet Staining for 30 minutes. Plates were scanned and quantification was performed in the ImageJ program with ColonyArea plugin (29).

### AnnexinV/7AAD assay

Apoptosis assessment was performed using Muse® Annexin V/Dead Cell kit (Luminex, MCH100105, USA) according to the manufacturer’s instructions. Conditioned medium and cells were collected and centrifuged at 1200 rpm for 5 minutes. Pellet was washed with 500 µl cold PBS containing 1%FBS followed by another centrifugation. Pellet was resuspended with 75 µl cold PBS containing 1%FBS and 75 µl reagent, and incubated for 20 minutes at dark, at RT. Measurement and analysis were performed with Muse® Cell Analyzer (Merck, Germany).

### Cell cycle analysis

Muse® Cell Cycle Assay kit (Luminex, MCH100106, USA) was used for cell cycle experiments according to the instructions. Cells were collected and centrifuged at 300xg for 5 minutes. Pellet was washed 2X with PBS. Cells were fixed as 1×10^6^ cells per 1 ml of 70% Ethanol for minimum 3 hours at −20°C. Ethanol was removed with centrifugation followed by wash with PBS. Pellet was resuspended with 150 µL of reagent and incubated at dark for 30 minutes at RT. Muse® Cell Analyzer (Merck, Germany) was used for measurements and analysis.

### Western blotting

The cell pellet was resuspended in NP-40 lysis buffer containing 1% NP-40, 50 mM Tris, 250 mM NaCl, 1X EDTA, 0.02% NaN3, 1 nM PMSF and 1X protease inhibitor cocktail, followed by incubation on ice for 30 minutes. The sample was centrifuged at maximum speed for 10 minutes at 4°C and supernatant was transferred for protein quantification. Pierce’s BCA Protein Assay Kit (Thermo Scientific, 23225, USA) was used to quantify isolated proteins. 20-30 µg of protein was boiled with 4X loading dye (4X Laemni buffer (Bio-rad, 1610747) with 1:10 beta-mercaptoethanol) at 95°C for 10 minutes. Samples were loaded to precast 4-12% Mini Protean TGX Precast Gel (Bio-rad, 456-1044). Gel was transferred onto PVDF membrane using Bio-RadTrans-Blot® Turbo™ Transfer System (Bio-Rad, USA). The membrane was blocked in 5% blocking grade blocker, or 5% Bovine Serum Albumin (BSA) prepared in 1X TBS-T for 1 hour at room temperature. Membrane was incubated with primary antibody overnight at 4°C. Next day, the membrane was washed 3x with 1X TBS-T and incubated with HRP conjugated secondary antibodies at RT for 1 hour. Visualization was performed using Li-Cor Odyssey® FC Imaging System. Primary and secondary antibodies used in the experiments were listed in **supplementary table 1.**

### Immunofluorescence and imaging

U87MG-RT cells were seeded on coverslips at PT8 following RBBP4 KO. At PT9, cells were fixed with 4% Paraformaldehyde and washed. Cells were permeabilized with 0.3% Triton-X in PBS for 5 minutes at RT followed by blocking with 5% BSA in PBS for 30 minutes at RT. Cells were incubated with the primary antibodies anti-α-tubulin (DM1A) (Sigma, T9026) 1:250 in 1% BSA and RBBP4 (RBAP48) (Proteintech, 20364-1-AP) 1:250 in 1% BSA overnight at 4 °C. The next day, cells were washed with PBS and incubated with Alexa Fluor 488 Anti-rabbit secondary antibody (Invitrogen, A11008) 1:250 and Alexa Fluor 594 Anti-mouse secondary antibody (Abcam, ab150116) 1:250 in 1% BSA in the dark for 2 hours at RT. Coverslips were mounted with VECTASHIELD® Antifade Mounting Medium (Vector Laboratories, USA) and visualized on Leica SP8 Laser Scanning Microscope (Germany). The images were analysed using CellProfiler to segment the nucleus with DAPI channel. Otsu thresholding method with three class thresholding was used to segment the cell boundaries. The signal analysis was performed inside the nuclear segmentation and the cell size analysis was performed by calculating the cell boundary mask area.

### Transcriptome analysis

High quality RNA was isolated with MN Nucleospin RNA isolation kit. U87MG and U87MG-TR samples were processed by BGI (Hong Kong, China); U373, U373-TR, U87MG-TR NT and U87MG-TR RBBP4 knock-out samples were processed at the University of Oxford (Oxford,UK), where library preparation and sequencing were performed. Analysis of the data was performed by the Bioinformatics Core in KUTTAM. Briefly, total RNA was DNase I-treated, cleaned, and concentrated using the Zymo RNA Clean & Concentrator kit (Zymo Research). Poly(A) mRNA was enriched using the NEBNext Poly(A) mRNA Magnetic Isolation Module (New England Biolabs, Ipswich, UK). Sequencing libraries were constructed with the NEBNext Ultra II RNA Library Prep Kit (New England Biolabs) following the manufacturer’s guidelines. RNA integrity was evaluated using High Sensitivity RNA ScreenTape on an Agilent 4200 TapeStation. Single-indexed, multiplexed libraries were sequenced on an Illumina NextSeq 500 system using a NextSeq 500 v2 kit (FC-404-2005; Illumina, San Diego, CA) for paired-end sequencing with a read length of 42 bp.

Sequencing reads were quality-assessed using FastQC, followed by alignment to the GENCODE GRCh38 human reference transcriptome using STAR (30). Transcript abundance was quantified using Salmon (31) based on the Transcripts Per Million (TPM) method. Transcript-level estimates were aggregated to the gene level using tximport (32). Differential gene expression analysis was performed using the DESeq2 package (33). For naïve-resistant sequencing pairs, genes with a log2 fold change (LFC) < –1 for downregulated or > 1 for upregulated were considered differentially expressed, while for NT1 vs RBBP4 knockout, thresholds of LFC < –0.5 and > 0.5 were applied. Gene Set Enrichment Analysis (34) was conducted on LFC-ranked gene lists using GSEA software, referencing gene sets from the MSigDB database. Overlap analysis was carried out using MSigDB gene sets to identify significantly enriched biological pathways.

### qRT-PCR

500-1000 ng of total RNA was used for cDNA synthesis using M-MLV Reverse Transcriptase kit (Invitrogen, USA). qRT-PCR reaction was performed using LightCycler 480 Instrument II (Roche, Switzerland) as triplicates per sample. Thermal Cycler reaction conditions were as follows: 95°C for 5 minutes as pre-heating step, 95°C for 10 seconds, 60°C for 30 seconds, 72°C for 30 seconds as 45 cycles. GAPDH was used for normalizations and calculations were performed by 2^-DeltaCT^ method. Primers used for qPCR are listed in **supplementary table 2**.

### Lentiviral packaging and transduction

For lentivirus and retrovirus production, 2.5×10^6^ HEK293T cells were seeded to 10 cm culture dishes. Next day, DNA mixture containing 2500 ng Vector, 2250 ng Gag-Pol (psPAX2 for lentivirus, pUMVC for retrovirus) and 225 ng VSVG was prepared in serum free DMEM. DNA mixture was added to Fugene mixture containing 15 µl FuGENE® 6 Transfection Reagent (Promega, USA) and 185 µl serum free DMEM and incubated for 30 minutes at RT. Later, mixture was added to the cells dropwise. Media were replaced with fresh media 16h later as 8 ml/plate. Supernatant containing viral particles was collected at 48- and 72-hour post-transfection and filtered through 45 µM filter (35). To concentrate, filtered supernatant was mixed with 5x PEG8000 (Sigma-Aldrich, USA) (dissolved in PBS as 50% (w/v)) for 1x final concentration, and kept in 4°C for 1-3 days. Supernatants were centrifuged at 2500 rpm for 20 minutes and virus pellet was resuspended in PBS as 100x (35). Viral titers were determined by seeding the cells as 1×10^5^ cells/well in 6-well plates. The following day, transduction at concentrations 1 µl, 10^-1^ µl, 10^-2^ µl and 10^-3^ µl was performed in the presence of 8 μg/ml protamine sulphate (Sigma-Aldrich, USA) overnight. Later, transduction media were replaced with fresh DMEM. Antibiotic selection, depending on the plasmid of interest, was initiated on the second day of transduction and completed once all uninfected cells were eliminated. Remaining cells in infected wells and unselected/uninfected wells were counted and compared to calculate viral titer.

### EPIKOL screening and analysis

For stable Cas9 expression, all cells were infected with LentiCas9-blast (Addgene # 52962) virus and selected with blasticidin. The pooled EPIKOL lentivirus targeting approximately 800 different genes was packaged as described. Cas9 expressing U87MG-TR and U373-TR cells were transduced with EPIKOL lentivirus (MOI:0.4) with 1000X coverage. Cas9 expressing NHA cells were transduced as 125X coverage with MOI:0.4. U87MG-TR and U373-TR screens were performed as triplicates, NHA screen was performed as a single replicate. Puromycin selection (1-2 µg/ml) was performed for 3-4 days, and first samples were collected after puromycin selection as 8×10^6^ cells for resistant cells and 1×10^6^ cells for NHA. Cells were passaged until Cas9 activity was complete (19-20 days for U87MG-TR, 17-18 days for U373-TR). Next, U87MG-TR and U373-TR cells were separated into two groups and treated with TMZ (200 µM and 250 µM of TMZ, respectively) or DMSO. Cells were passaged until both groups reach 14-15 population doublings and endpoint pellets were collected as 8×10^6^ cells. NHA cells were passaged for 6 weeks post-transduction and endpoint pellets were collected as 125X. Genomic DNA (gDNA) isolation was performed using MN Nucleospin Tissue kit according to the manual. Isolated gDNA served as the template for library preparation of samples for Next Generation Sequencing (NGS) (Illumina). Total of 13.2 µg DNA was amplified per sample to achieve 250X gRNA coverage from initial PCR reaction. Illumina adapters (stagger and index sequences) were added with the second PCR reaction. Thermal cycler conditions for PCRs were as follows: 95°C for 3 minutes, 95°C for 25 seconds, 65°C for 20 seconds and 72°C for 15 seconds (17X for external PCR and 23X for internal PCR) and 72°C for 3 minutes. PCR products were loaded on 2% Agarose gel and isolated using MN Nucleospin Gel and PCR Cleanup kit. Samples were sequenced by Genewiz (NJ, USA) as at least 10 million reads/sample. Analysis of EPIKOL CRISPR screening data was performed with Model-based Analysis of Genome-wide CRISPR-Cas9 Knockout (MAGeCK) tool v0.5.9 (36) by grouping samples by median normalisation, variance modelling, robust rank aggregation (RRA) and group specific essential gene identification. Downstream analyses were performed with a combination of tools, as we previously described (23).

### Live cell imaging

Cas9 expressing U87MG-TR cells transduced with RBBP4 g1, g2 or NT1 were infected with PGK-H2BmCherry (Addgene #21217) lentivirus at PT6. The next day, cells were seeded to 6-well plates as duplicates with 150.000 cells/well density. Between PT8-PT10, 48h live cell imaging was performed with Cytation 5 (BioTek, USA). Phase contrast and Texas Red fluorescent-based images were taken as 3×3 fields/well every 30 minutes. Gen5 software (BioTek, USA) was used to count mCherry + cells at each time point from images. Mitotic time was calculated by tracking individual cells through mitosis in ImageJ and registering the number of frames during which the cell divided and multiplying it with 30 minutes. Representative videos of the assay for NT1, RBBP4 g1 and g2 can be found in **supplementary videos**.

### GFP competition assay with flow cytometry

Cas9 expressing U87MG-TR and U373-TR cells were infected with pBabe-Hygro-GFP (Addgene #61215) and selected with 150 µg/ml Hygromycin. Only Cas9 expressing cells were transduced with NT1 while Cas9-GFP expressing cells were transduced with RBBP4 g1, g2, RPL9 or NT1. To determine the optimum time required for Cas9 activity, Cas9-GFP expressing cells were transduced with T1 (GFP-targeting gRNA) or NT1. Puromycin selection was performed as usual. Cas9 only and GFP only cells were maintained as controls. At PT5, Cas9 and Cas9-GFP expressing cells were mixed 1:1 ratio where half of the population was seeded to 24-well plate as duplicates and other half was used for Day0 measurement. Flow cytometry samples were washed 2X with PBS containing 1% FBS, GFP readouts were taken using Cytoflex Flow Cytometer (Beckman Coulter, USA) every 4 days.

### In vivo experiments

All animal experiments were approved by the institutional ethical committee of Koç University (2016–15). U87MG and U87MG-TR cells were infected with firefly luciferase-mCherry (FLUC-mCherry) viruses as described (24). Tumor cells (100,000 cells/mice) were implanted to 6-8-week-old non-obese diabetic/severe combined immunodeficiency (NOD/SCID) mice orthotopically with stereotaxic tools from bregma, AP: −2 mm, ML:1.5 mm, V (from dura):2 mm). Visualization of tumors was performed with IVIS Lumina III (Perkin Elmer, USA) under isoflurane anesthesia via intraperitoneal injection of D-Luciferin (150 µg/g body weight) at different time points. Mice bearing naïve or resistant tumors were further divided into two groups (n=5/group) and TMZ or DMSO was administered as 2 mg/kg in PBS (37). On day 24, brain samples were collected from mice and fixed in 10% neutral buffered formalin (NBF).

### Immunohistochemistry

Sectioning, H&E and Ki-67 staining were performed at Koç University Hospital, Pathology Department. Mice brains were sectioned in coronal slices from anterior to posterior and tissue processing was performed using tissue processor (Tissue-Tek CIP 6 AI, Sakura, Japan). Tissues were dehydrated by sequential immersion in graded ethanol solutions (70%, 95% and 100%) followed by xylene treatment and paraffin embedding. 5 µm thick slices were generated from paraffin-embedded tissues with a microtome. For hematoxylin and eosin (H&E) staining, tissue sections were deparaffinized via sequential xylene and rehydration using graded alcohol washes. Sections were stained with H&E staining according to Leica protocol (38). Ki-67 staining was performed on Ventana XT Autostainer using Ki67 antibody (Rabbit monoclonal, Clone: SP6, dilution 1:100, Biocare Medical, CA, USA). Slides were scanned with Philips Ultra-Fast Scanner (UFS) and evaluated with Philips Image Management System. QuPath was used to select and evaluate five areas with high Ki-67 staining and positive cell detection algorithm was performed after calibration (the cell size, pixel height, and width in microns).

### Statistical Analysis

Statistical analysis of the experiments was performed using GraphPad Prism 10.0. Two groups were compared using student’s t-test, more than two groups were compared using two-way ANOVA. * p<0.05, ** p<0.01, *** p<0.001.

### Data Availability

EPIKOL screening and RNA sequencing data are deposited to NCBI GEO database with the accession numbers GSE286427 and GSE286225, respectively.

## RESULTS

### Temozolomide-resistant glioblastoma cells can be generated with long term Temozolomide exposure and maintained in vivo

To investigate the mechanisms of chemoresistance in glioblastoma, we generated *in vitro* Temozolomide-resistant models (**Supp. Fig. 1A**). We utilized dose escalation method for U87MG cell line and single dose treatment for U373 cell line. Starting dose of 5 µM TMZ was administered to U87MG cells and dose was increased until 200 µM, where cells exhibit morphological changes. 250 µM of TMZ was administered to U373 cells for 15 consecutive days followed by recovery and maintained in 250 µM of TMZ for two months. Generated models were named as U87MG-TR and U373-TR, respectively. Both models displayed increased IC_50_ values compared to their parental cell lines (**Fig 1A**). Long term colony formation assay results demonstrated that resistant cells maintain the ability to form colonies under high TMZ concentrations whereas parental cells fail to grow after 10 µM of TMZ (**Fig. 1B, 1C, Supp. Fig. 1B, 1C**). Our cell cycle analysis showed significant increase in the percentage of G2/M population upon 72h TMZ treatment for parental cells, whereas no change was observed for the resistant cell lines (**Fig. 1D, Supp. Fig. 1D**). To test the resistance phenotype *in vivo*, we implanted Fluc-mCherry labelled U87MG and U87MG-TR cells intracranially to NOD-SCID mice (n=5/group), and measured tumor growth over 24 days with bioluminescence imaging (**Fig. 1E**). Once the tumor growth was observed on Day8 after implantation, the mice were divided into two groups, ensuring the tumor volumes were similar in each group. One group received TMZ (2 mg/kg) for 5 consecutive days and the other group served as controls. On Day24, mice were sacrificed for histology. Representative images for Day0 and Day24 show the increase/decrease in the Luciferin signal for each group, demonstrating the loss in tumor volumes in the U87MG tumors upon TMZ treatment (**Fig. 1F**). On the contrary, tumors formed from U87MG-TR cell line did not show any decrease in signal upon TMZ treatment, indicating the maintenance of resistance *in vivo*. Notably, the growth rates of U87MG-TR tumors were significantly higher than naïve U87MG cell-derived tumors. Histopathological examination of brain sections demonstrated the presence of U87MG tumors with compact borders in the untreated group, whereas no tumor was observed at the TMZ-treated group besides remaining necrotic tissues (**Fig. 1G**). Interestingly, very large and invasive tumors were observed in U87MG-TR groups regardless of TMZ administration, indicating the aggressive nature of the TMZ-resistant cells. Assessment of tumor cell proliferation with Ki67 staining revealed significantly higher proliferation index tumors derived from U87MG-TR cells compared to those from U87MG parental cells, independent of TMZ treatment (**Fig. 1H**). These results provide evidence to the successful generation of Temozolomide-resistant models from U87MG and U373 cell lines and demonstrate their ability to grow under high TMZ concentrations. One of these models, U87MG-TR, can form highly aggressive and proliferative tumors *in vivo*, serving as a potent model of therapy-refractory recurrent glioblastoma.

**Figure 1:**
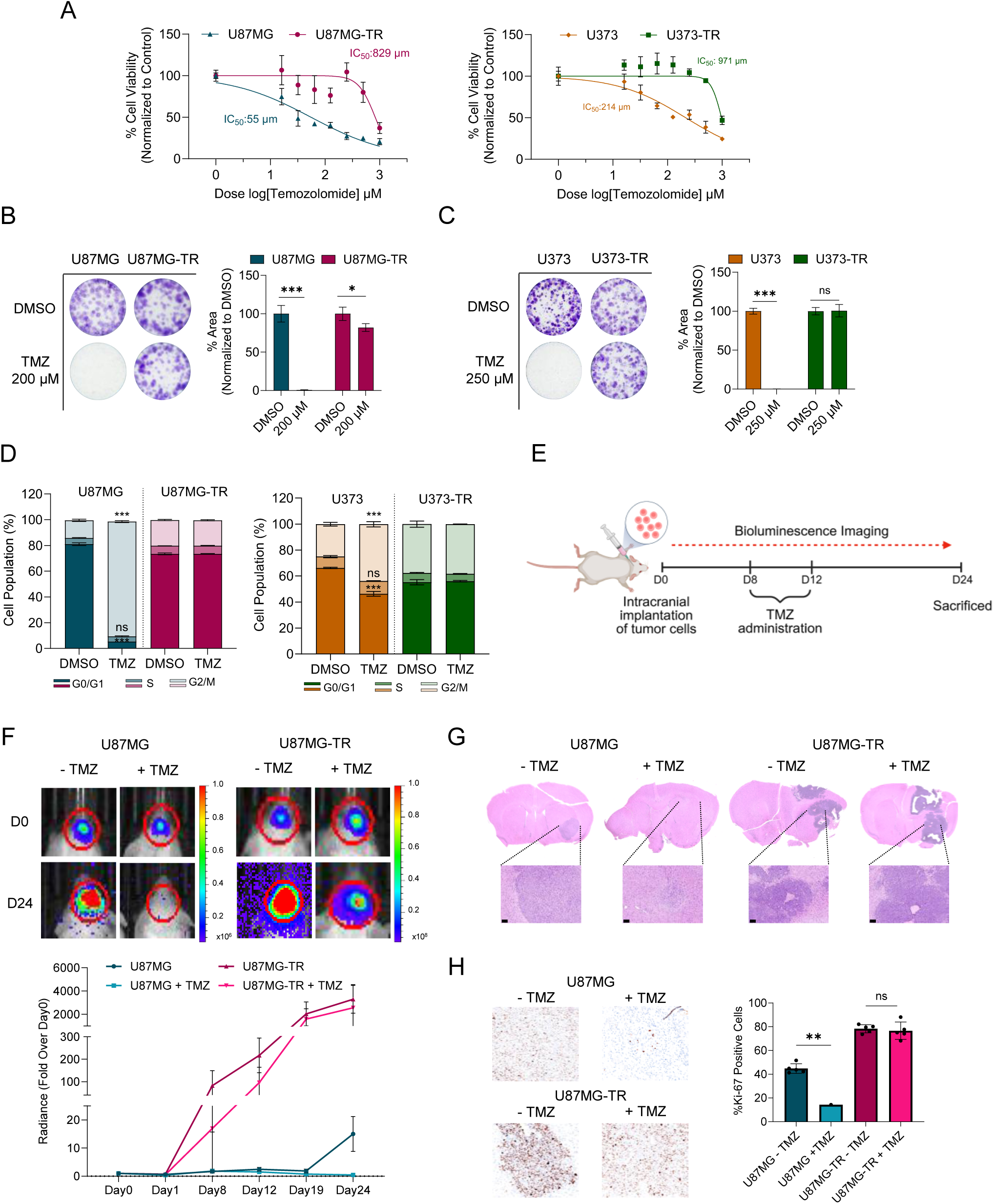
Generation and characterization of Temozolomide-resistant cell lines. **A.** MTT cell viability results showing IC_50_ values of parental and Temozolomide-resistant models. **B, C.** Long term colony formation assay results upon TMZ administration. Quantification of colony results was performed by ImageJ colony area plugin, normalized to DMSO of each cell line. P values were determined by t-test in comparison to DMSO of each cell line; *p < 0.05, **p < 0.01, ***p < 0.001. **D.** Cell cycle analysis after 72h TMZ treatment (200 µM for U87MG and U87MG-TR, 250 µm for U373 and U373-TR). P values were determined by Two-way Anova between DMSO-TMZ groups; *p < 0.05, **p < 0.01, ***p < 0.001. **E.** Schematics of intracranial implantation of U87MG and U87MG-TR cell lines. Figure generated with BioRender.com **F.** Representative images of Day0 and Day24 for experimental groups. Color scale for radiance (p/sec/cm^2^/sr): ×10^6^ for U87MG +/− TMZ Day0/24, U87MG-R +/− TMZ Day0, ×10^8^ for U87MG-R +/− TMZ Day24. Graph shows the radiance of tumors over 24 days. **G.** H&E staining of brain sections collected on Day24. Closeup scale bar :100 µm (10X). **H.** Ki-67 staining showing the proliferative cells in tumor sections (20X). Graph shows the % of Ki-67 positive cells over all cells in the section as 5 stained areas. P values were determined by t-test; *p < 0.05, **p < 0.01, ***p < 0.001.

### Transcriptome analysis demonstrates signature genes and dependency to MGMT

To investigate the molecular alterations that were acquired in TMZ-resistant cells, we performed transcriptome analysis and assessed gene expression patterns in naïve and resistant cell lines. In U87MG-TR cells, 1121 upregulated and 988 downregulated, in U373-TR cells, 1389 upregulated and 788 downregulated genes were identified as demonstrated in Volcano plots with LFC<-1, >1 filters (**Fig. 2A).** Total of 182 genes were commonly upregulated in U87MG-TR and U373-TR cells (**Fig. 2B**). Gene Ontology (GO) analysis using the Molecular Signature Database (MsigDB) revealed that these 182 genes are associated with processes involved in neurogenesis, cell movement and cell surface modifications (**Supp. Fig. 2A**). Among the 182 commonly upregulated genes, the top 15 *were MGMT, TUBA3C, ROBO2, RNF43, MTUS1, COLEC10, SEMA6D, PASD1, CES1, CSMD1, TMEM132D, CELF2, SLC14A1, KLHL4* and *ESM1*. qRT-PCR analysis validated the upregulation of these genes in U87MG-TR cell line compared to U87MG (**Supp. Fig. 2B**). Role of MGMT in TMZ response and resistance is well established (11). We confirmed its overexpression at the protein level using western blotting in U87MG-TR and U373-TR cell lines, as opposed to undetectable protein expression in the naïve U87MG and U373 cell lines (**Fig. 2C**). To further investigate the function of MGMT in our resistant models, we used CRISPR/Cas9 to knock-out (KO) MGMT with two independent gRNAs. Both gRNAs successfully reduced MGMT expression in U87MG-TR and U373-TR cell lines compared to non-targeting control (NT1) (**Fig. 2D).** Long term colony formation assays showed that MGMT KO cells grew similarly to controls in the absence of TMZ administration. However, growth of U87MG-TR and U373-TR cell lines was significantly reduced upon TMZ administration (**Fig. 2E, 2F**). Overall, these results indicate that TMZ-resistant cells exhibit marked changes in gene expression with MGMT being one of the top upregulated genes in both models. Knock-out of MGMT reverses the resistance in U87MG-TR and U373-TR, suggesting a strong dependence on MGMT for the resistant phenotype.

**Figure 2:**
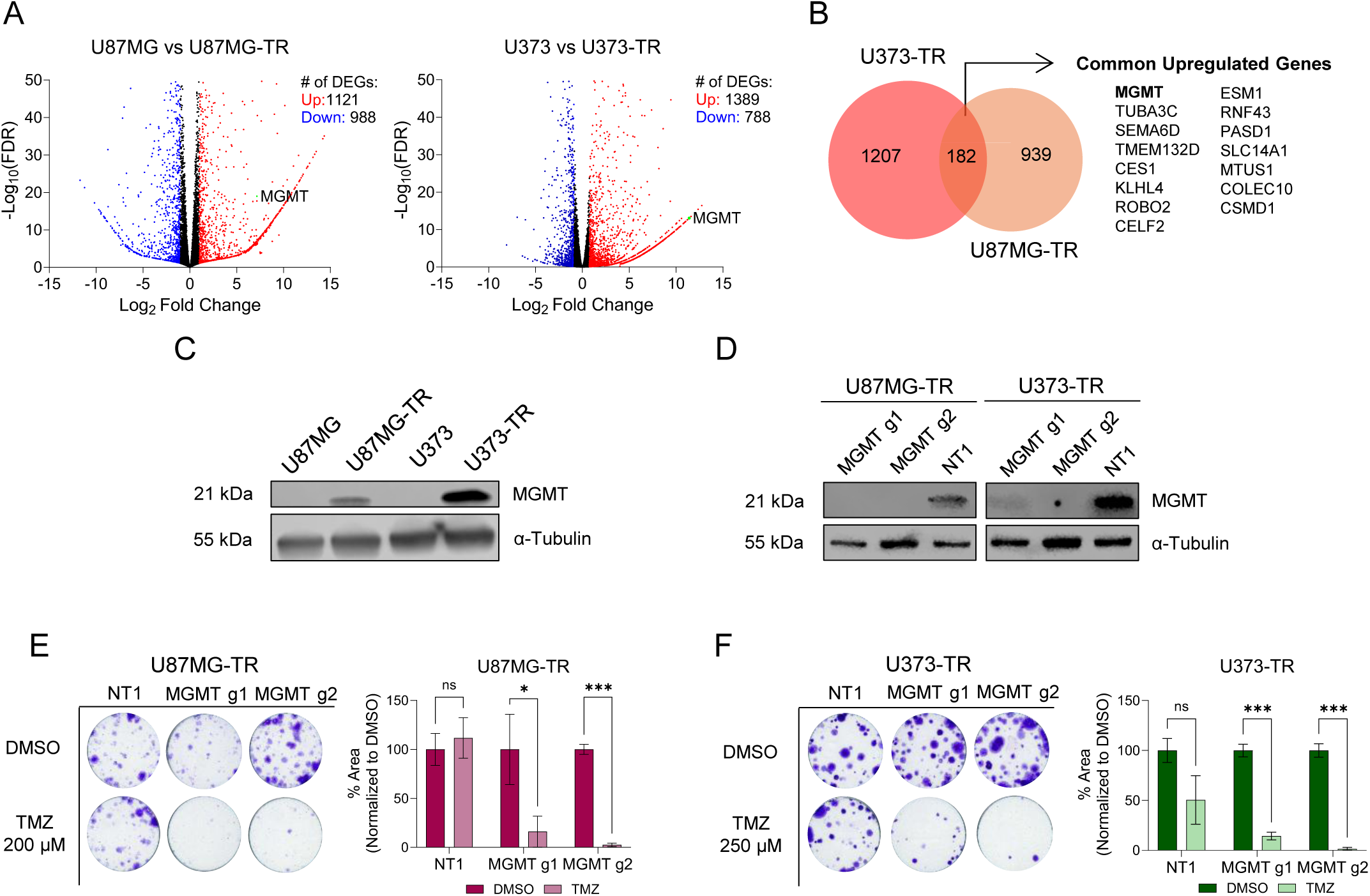
Acquired TMZ resistance is associated with high MGMT expression. **A.** Volcano plots showing upregulated (red) and downregulated (blue) differentially expressed genes between parental-resistant cell lines. Cutoff LFC<-1, LFC>1. **B.** Venn diagram of common upregulated genes in both models. The top 15 genes are listed **C**. MGMT protein expressions shown by western blot in naive and resistant cell lines. **D.** MGMT Knock-out validation by western blot using two different gRNAs in U87MG-TR and U373-TR. **E, F**. Long term colony formation assay results for MGMT KO with TMZ administration. Graphs showing the quantification of colonies by ImageJ colony area plugin. P values were determined by t-test in comparison to DMSO of each cell line; *p < 0.05, **p < 0.01, ***p < 0.001.

### Epigenetic Knock Out Library (EPIKOL) screens reveal potential TMZ sensitizers

To identify potential epigenetic mechanisms that can overcome therapy resistance, we utilized chromatin-focused CRISPR/Cas9 screening using our previously generated library, Epigenetic Knock Out Library (EPIKOL) (23). EPIKOL contains 744 epigenetic modifiers along with 35 essential genes as 10 gRNAs/gene and 80 non-targeting gRNAs with the total of 7870 gRNAs. First, the time frame where Cas9 enzyme successfully knock-out gene of interest was calculated. We used GFP-targeting gRNA (T1) in Cas9 and GFP stable cell lines, mixed with Cas9-only stable cell lines infected with NT1 gRNA (**Supp. Fig. 3A**). The results showed decrease in %GFP levels beginning on Day 4 (PT9), which stabilized by Day 12 (PT18) (**Supp. Fig. 3B**). Based on these findings, Cas9 activity for U87MG-TR and U373-TR cells were determined to be around PT18. Next, we performed EPIKOL screens with U87MG-TR and U373-TR and administered TMZ or DMSO following puromycin selection and completion of Cas9 activity (**Supp. Fig. 4A**). To find sensitizers, we compared the endpoints of the groups as TMZ and DMSO with LogFoldChange (LFC)<0 and p-value<0.05 cutoffs. By comparing the results from both screens, we identified four potential sensitizing genes: *MGMT*, *APEX1* (Apurinic/Apyrimidinic Endodeoxyribonuclease 1), *PCGF6* (Polycomb Group Ring Finger 6) and *SET* (SET Nuclear Proto-Oncogene) (**Supp. Fig. 4B**). LFC graphs showed that all four were significantly depleted during the screens, with *MGMT* being one of the top depleted genes (**Supp. Fig. 4C**). To validate our findings, we performed long term colony formation assay upon knockout of four candidate genes along with a positive control (RPL9) and negative control (NT1) in the presence of TMZ or DMSO. Surprisingly, the results showed that only MGMT KO significantly sensitized resistant cells to TMZ, whereas the loss of SET, APEX1 and PCGF6 showed minimal effects (**Supp. Fig. 4D, 4E**). qRT-PCR confirmed the successful downregulation of MGMT, APEX1, SET, PCGF6 and RPL9 levels (**Supp. Fig. 4F**). These results demonstrate the effectiveness of the unbiased EPIKOL screen in identifying MGMT as the principal regulator of TMZ response, highlighting it as a primary intervention target to overcome TMZ resistance in our models.

### EPIKOL screens identify selective epigenetic vulnerabilities of TMZ-resistant cell lines

Although the identification of MGMT dependency in acquired TMZ resistance was of interest, we sought to further investigate the epigenetic vulnerabilities of the cells in their TMZ-resistant state, specifically in the absence of TMZ treatment. We analyzed InitialPoint vs EndPoint (AP vs DMSO) comparisons of EPIKOL screens (**Fig. 3A**) to plot the gRNAs that were depleted over time. gRNA count distribution graphs were generated to determine gRNA abundance using read per million normalizations of raw counts followed by Log2 conversion (**Fig 3B**). Accordingly, while the positive control essential genes were depleted in both TMZ-resistant cell lines, non-targeting gRNAs showed either no or minimal changes, highlighting the success of our screens. Separately, we performed an EPIKOL screen with Normal Human Astrocytes (NHA) to identify essential genes for non-malignant cells and compare with cancer cell lines to ultimately identify cancer specific vulnerabilities (**Supp. Fig. 5A, 5B**). By excluding the genes identified in NHA screen and positive control essential genes, we identified seven epigenetic vulnerabilities in TMZ-resistant cells: Retinoblastoma Binding Protein 4 (*RBBP4*), Proliferating Cell Nuclear Antigen (*PCNA*), Exosome Component 2 (*EXOSC2*), Cyclin Dependent Kinase 2 (*CDK2*), BRCA1 Associated Deubiquitinase 1 *(BAP1*), NSL1 Component of MIS12 Kinetochore Complex (*NSL1*) and DNA Topoisomerase II Alpha (*TOP2A*). LFC graphs demonstrated the depletion of these genes in U87MG-TR and U373-TR cells (**Fig. 3D**), but not in the NHAs (**Supp. Fig. 5C**). Gene specific gRNA counts showed consistent depletion at the endpoint in TMZ-resistant cell lines with modest or no effect in NHA cells (**Fig. 3E, Supp. Fig. 5D**). To validate the effect of these genes on U87MG-TR and U373-TR cell lines, we cloned individual gRNAs and performed long term colony formation assay. The loss of all these hit genes significantly reduced the colony forming ability of U87MG-TR and U373-TR, with CDK2 being the exception for U373-TR (**Fig. 3E**). qRT-PCR confirmed significant depletion of mRNA levels for all target genes with the CRISPR approach (**Supp. Fig. 5E**).

**Figure 3:**
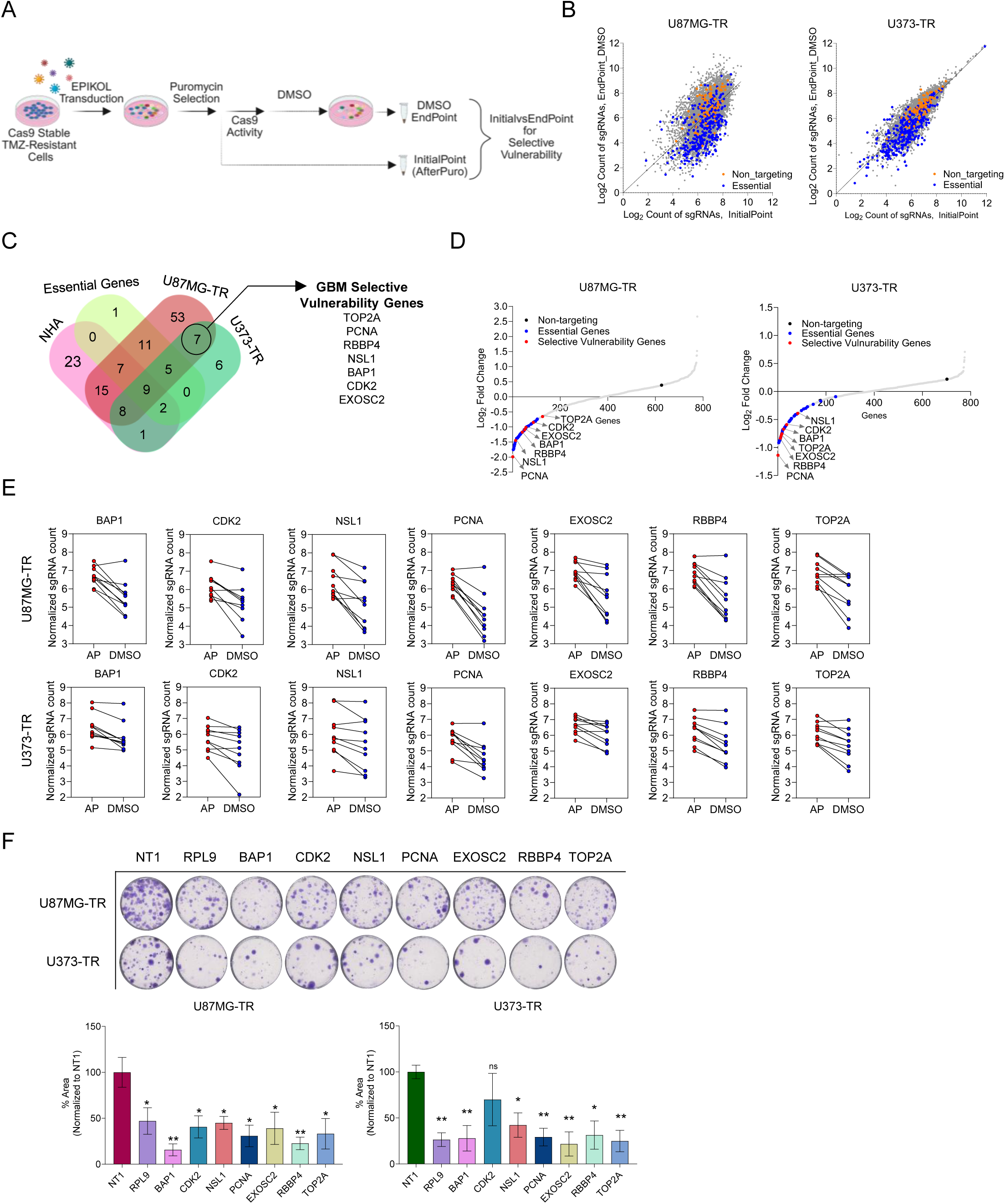
Multiple CRISPR screens identify epigenetic vulnerabilities of TMZ-resistant GBM cells. **A**. Schematic for EPIKOL screen strategy for U87MG-TR and U373-TR. Cells were infected with EPIKOL lentiviruses as 1000X coverage with MOI:0.4 and selected with puromycin. Initial pellet was obtained after puromycin selection (AfterPuro). Following Cas9 activity, cells were divided into two groups as DMSO and TMZ. Screen continued until each group reached 14-16 population doublings and endpoint pellets were collected. InitialPoint (AfterPuro) and DMSO-Endpoints were compared. Figure generated with BioRender.com **B.** gRNA distribution of EPIKOL screen results for resistant cell lines. **C.** Venn diagram showing number of common genes depleted in both resistant models, excluding hits from Normal Human Astrocyte (NHA) screen and essential genes within the library. **D.** Log2 Fold Change graphs depicting the vulnerability genes based on LFC scores. **E**. sgRNA count comparisons between InitialPoint (AfterPuro (AP)) and EndPoint (DMSO) for vulnerability genes. **F.** Long term colony formation assay for selective vulnerability genes. The quantification was performed with ImageJ Colony Area plugin, normalized to NT1. P values were determined by t-test compared to NT1; *p < 0.05, **p < 0.01, ***p < 0.001.

Together, these results revealed novel epigenetic vulnerabilities of TMZ-resistant cells, providing potential targets for therapeutic intervention.

### RBBP4 loss leads to decreased viability and leads to apoptosis

Among the identified genes, RBBP4 has recently been associated with TMZ sensitivity and cell survival through its role in DNA repair pathways (26–28). However, its specific involvement in acquired TMZ resistance remains unexplored. To investigate the function of RBBP4 in TMZ resistance models, we employed two independent gRNAs for RBBP4 and confirmed the significant reduction of protein expression in KO samples with western blotting (**Fig. 4A**). Colony formation assay conducted with RBBP4 gRNAs showed significant reduction in viability of cells comparable to positive control RPL9, in accordance with our previous observations (**Fig. 4B**, **Fig. 3F**). We then performed GFP competition assay to assess the contribution of RBBP4 to the viability of TMZ-resistant cells in a heterogeneous model (**Fig. 4C**). Briefly, GFP tagged Cas9 stable cells were transduced with gRNAs for RBBP4, RPL9 or NT1, whereas Cas9-only cells were transduced with NT1 alone. Cas9-only and GFP cells were mixed as 1:1 ratio and the percentage of GFP positive population was measured with flow cytometry every five days starting at PT5. GFP levels of RBBP4 KO + NT1 cells dropped below 25% in U87MGT-R cells and to 30% in U373-TR cells (**Fig. 4D**). Negative control groups (NT1+NT1) remained at 50% throughout the experiment for both cell lines, while positive control groups (RPL9+NT1) showed the greatest depletion. An MTT based growth curve assay further supported these findings, with RBBP4 KO cells exhibiting significantly reduced growth by Day 5 in both the U87MG-TR and U373-TR cells (**Fig. 4E**). In accordance with the reductions in growth, both the early and late apoptosis was induced in RBBP4 KO cells compared to NT1, as gauged by using Annexin/Dead Cell assay at PT12, suggesting that loss of RBBP4 led to a significant induction of apoptosis (**Fig. 4F**).

**Figure 4:**
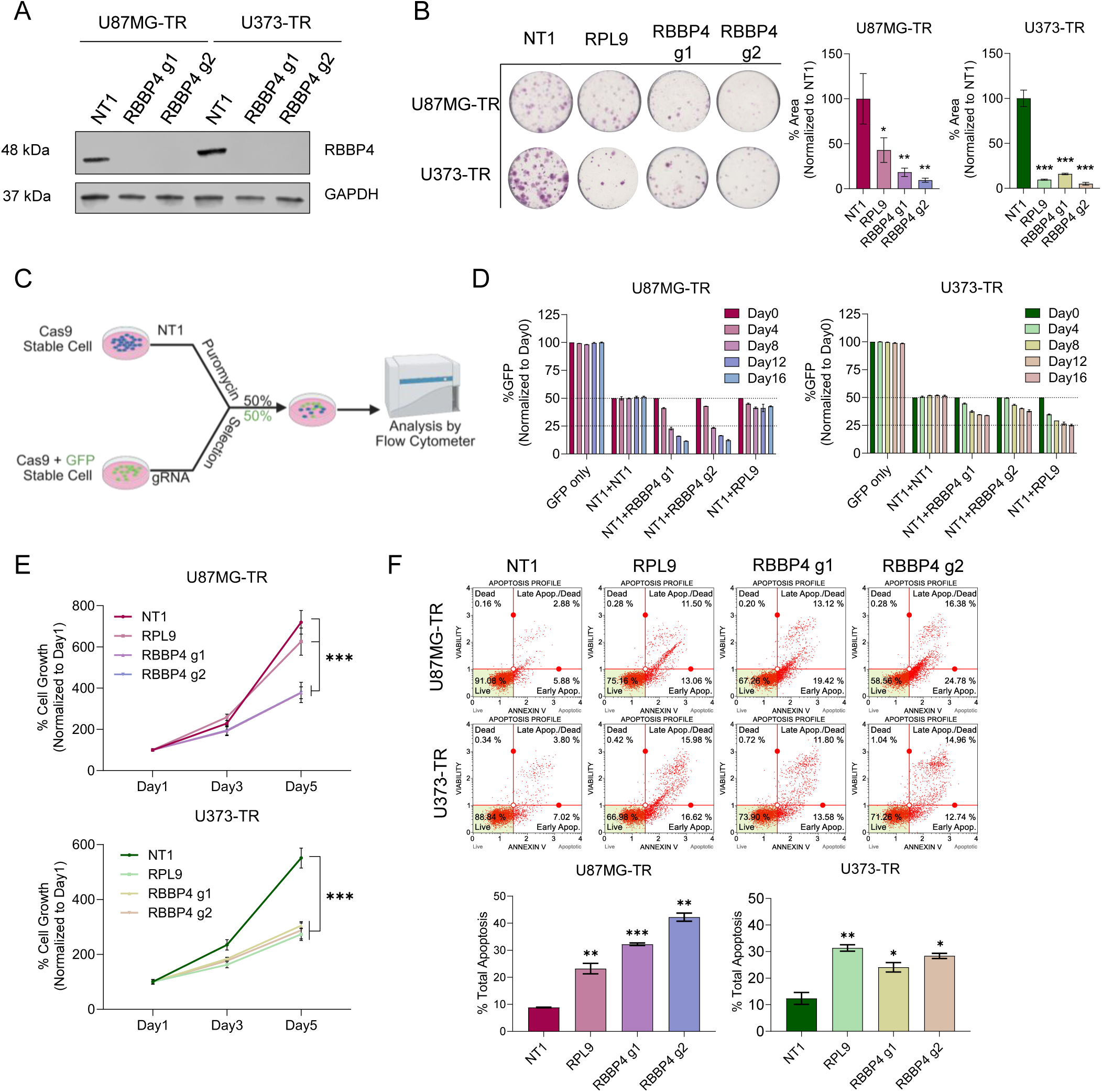
RBBP4 regulates survival of TMZ-resistant cell lines. **A.** Knock-out of RBBP4 with two guide RNAs was validated with western blot at post-transduction day 9 (PT9). **B.** Colony formation assay shows a decrease in viability upon KO. Quantification of colonies was performed with ImageJ colony area plugin. P values were determined by t-test compared to NT1; *p < 0.05, **p < 0.01, ***p < 0.001. **C.** GFP Competition assay. Cas9 only cells were infected with NT1 whereas Cas9+GFP cells were infected with gRNAs. Following puromycin selection, cells were mixed 50%-50% and GFP was measured every four days with Flow Cytometer. Figure generated with BioRender.com **D.** GFP competition assay results for U87MG-TR and U373-TR, normalized to Day0 (Posttransduction Day 5). **E.** Cell proliferation assessment via MTT viability assay at days 1, 3 and 5 (PT10, 12, 14 respectively). **F.** Annexin/Cell death assay results at PT12. Graphs indicate total apoptosis levels in both cell lines. P values were determined by t-test compared to NT1; *p < 0.05, **p < 0.01, ***p < 0.001.

To determine if the dramatic changes in cell proliferation and growth were exclusive to RBBP4 or shared with its homolog RBBP7(39), we knocked out RBBP7 in U87MG-TR and U373-TR cells. Colony formation assays showed that RBBP7 KO did not cause any significant reduction in U87MG-TR and led to only a slight decrease in U373-TR, with no significant decrease in cell viability (**Supp. Fig. 6A-B**), despite significantly decreased RBBP7 mRNA levels (**Supp. Fig. 6C**). To conclude, the phenotype observed upon RBBP4 KO was not phenocopied by its homolog RBBP7, attesting to the specific function of RBBP4 in the acquired TMZ-resistant cell state.

To then test the effectiveness of RBBP4 loss in naïve glioblastoma cells, that were either innately TMZ-sensitive or TMZ-resistant, we knocked-out RBBP4 in U87MG and T98G cell lines, respectively, in parallel with the U87MG-TR cell line (**Fig. 5A, B, C**). Long term colony formation assays with TMZ (U87MG: 10 µM, U87MG-TR: 200 µM, T98G: 125 µM) or DMSO treatments showed that RBBP4 KO decreased colony forming ability in all three cell lines, but TMZ sensitization was observed only in U87MG cells (**Fig. 5D**). Western blotting confirmed the loss of RBBP4 protein across all knockout cell lines (**Fig. 5E**). To test whether RBBP4 function would affect MGMT expression, as previously suggested (26,27), we examined MGMT expression levels in acquired and innately TMZ-resistant U87MG-TR and T98G cells, respectively, following RBBP4 KO. In U87MG-TR no changes were observed in MGMT mRNA or protein levels, whereas T98G cells exhibited a significant decrease in both mRNA and protein levels, in accordance with previous observations (26) (**Fig. 5E**, **Fig. 5F**). These findings indicate that the phenotype caused by RBBP4 loss is not mediated by MGMT in our acquired TMZ-resistant model. To explore RBBP4 function further, we tested its relationship with MRN complex (MRE11, RAD50 and NBN (NBS1)), as previously (28). We checked mRNA levels of MRN complex members for U87MG-TR and T98G after RBBP4 KO, however no significant change was observed for both cell lines (**Fig. 5G**). Overall, these results suggest that the mechanism behind RBBP4 loss-mediated reduction in cell viability is different than previously described pathways, highlighting a novel role for RBBP4 in acquired TMZ resistance.

**Figure 5:**
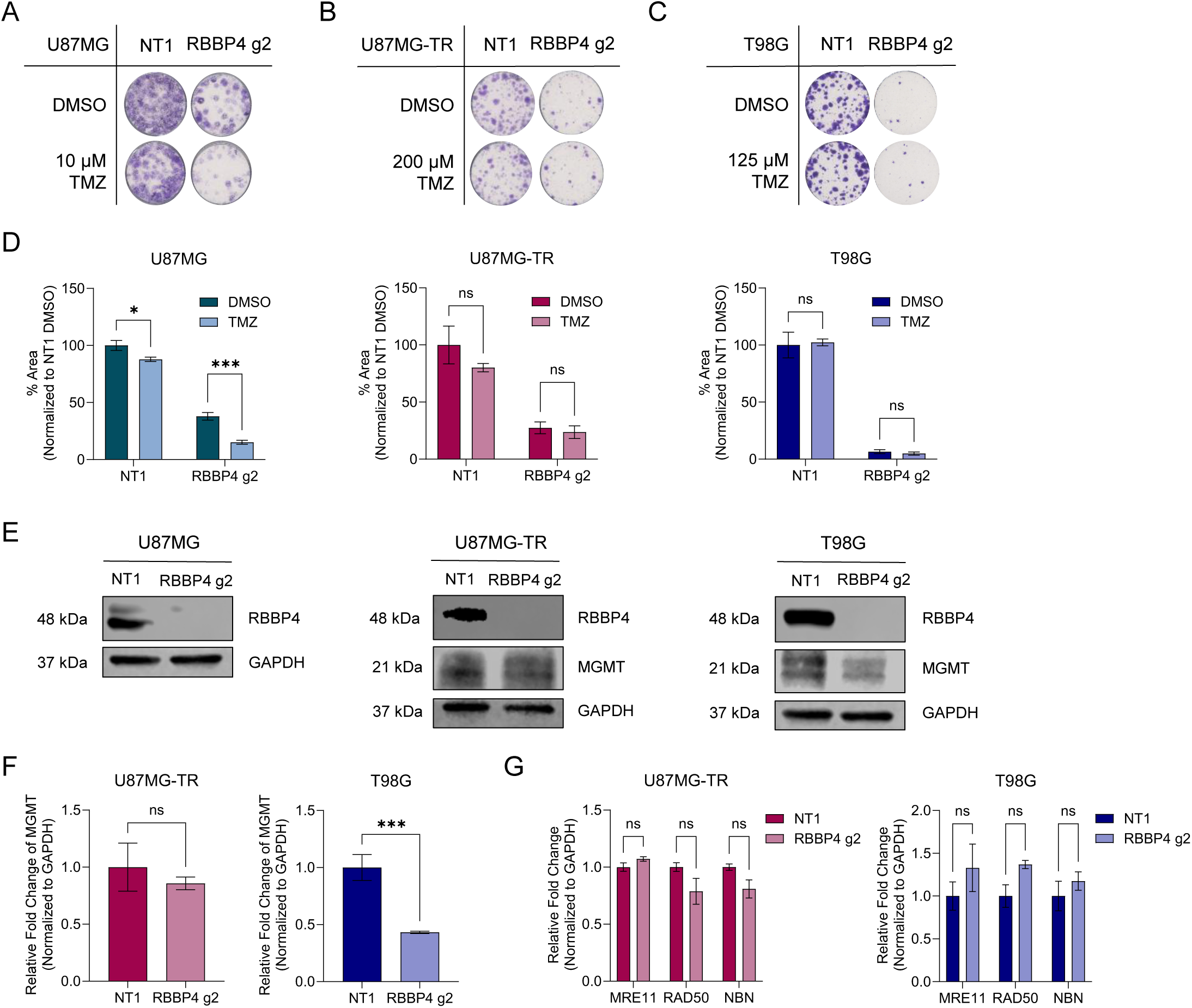
Effect of RBBP4 on naive, innately TMZ-resistant and acquired TMZ-resistant cell lines. **A, B, C.** Colony formation assay for U87MG, U87MG-TR and T98G showing long term viability upon KO with TMZ/DMSO. **D**. Quantification of colonies was performed by ImageJ program with ColonyArea plugin. P values were determined by t-test between DMSO-TMZ; *p < 0.05, **p < 0.01, ***p < 0.001. **E.** Western blot analysis of RBBP4 and MGMT for U87MG, U87MG-TR and T98G. F. RT-qPCR results for MGMT expression in U87MG-TR and T98G cell lines. P values were determined by t-test compared to NT1; *p < 0.05, **p < 0.01, ***p < 0.001. **G.** RT-qPCR results for MRE11-RAD50-NBN (MRN) complex members in U87MG-TR and T98G. P values were determined by t-test compared to NT1; *p < 0.05, **p < 0.01, ***p < 0.001.

### Transcriptomic changes upon RBBP4 depletion reveals cell cycle dependency

To understand the mechanism behind the RBBP4 phenotype, we performed RNA sequencing with U87MG-TR using both gRNAs for RBBP4 at PT9. Volcano plots show the DEGs and number of upregulated and downregulated genes with LFC<-0.5, >0.5 cutoffs (**Fig. 6A**). RBBP4 was one of the top downregulated genes, as a testament to the successful reduction of its expression upon KO. 321 downregulated, 756 upregulated and 220 downregulated, 665 upregulated genes were identified with RBBP4 g1 and g2, respectively. We focused on the 169 commonly downregulated genes shared between both gRNAs (**Fig. 6B**). Overlap analysis using the Molecular Signature Database (MsigDB) revealed significant enrichment for G2M checkpoint, E2F targets and mitotic spindle pathways among these shared downregulated genes (**Fig. 6C**). Gene Set Enrichment Analysis (GSEA) further confirmed the G2/M checkpoint as a significantly downregulated pathway upon RBBP4 loss (**Fig. 6D**). We generated a heatmap based on Log_2_ Fold Change values of genes found in all three, two or only one of these pathways (**Fig. 6E**). Next, we selected 10 genes from this list and validated RNA sequencing results using qRT-PCR (**Fig. 6F**). To conclude, transcriptome analysis showed that cell cycle related pathways was disrupted upon RBBP4 KO, leading to decreased expression of cell cycle-related genes.

**Figure 6:**
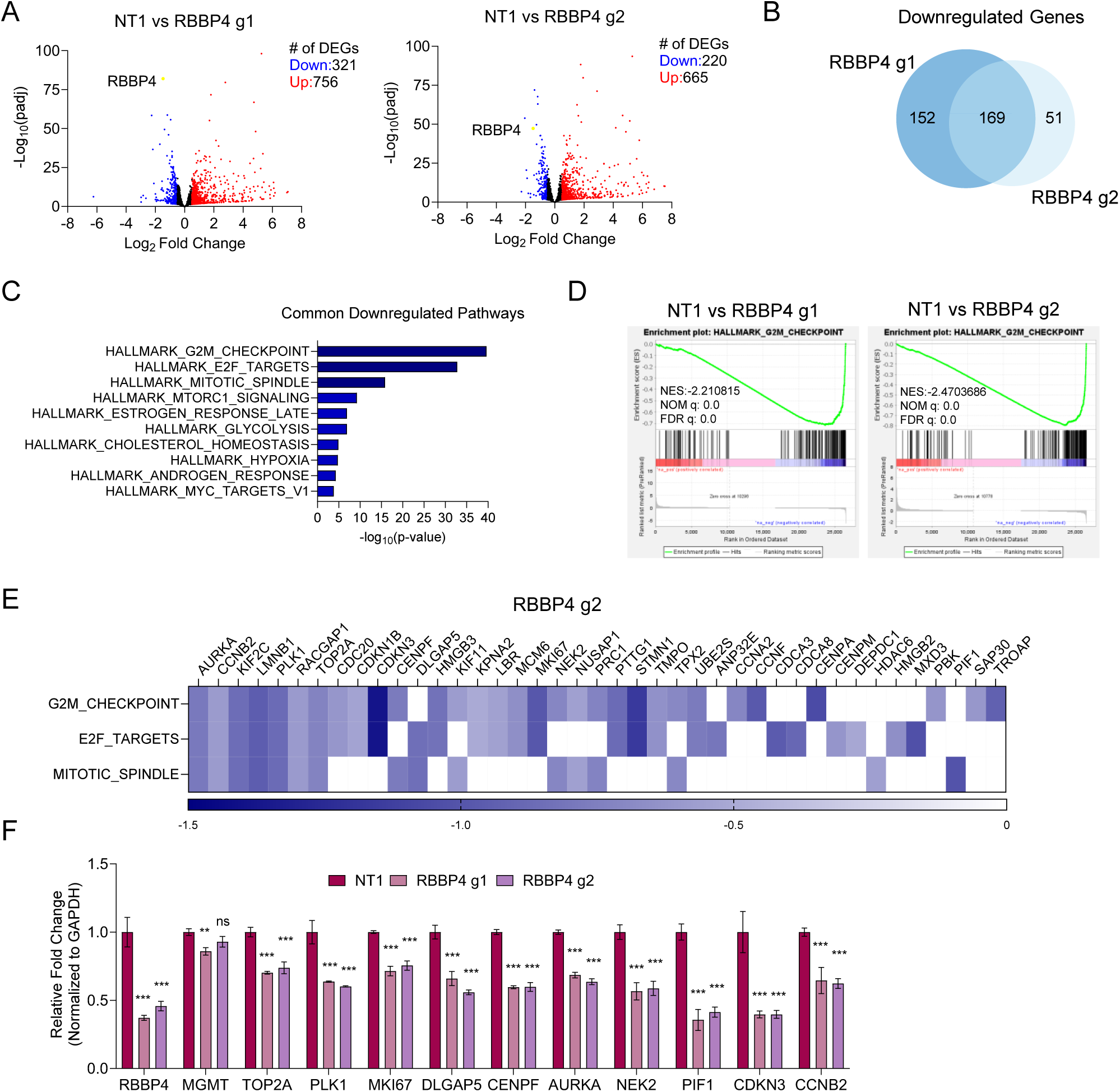
Transcriptome Analysis of RBBP4 KO in U87MG-TR cell line. **A.** Volcano plots showing the distribution of downregulated and upregulated genes with both gRNAs. LFC>0.5 and LFC<-0.5 cutoff were used. **B**. Venn diagram with the number of common downregulated genes in both gRNAs with LFC<-0.5 cutoff. **C**. Overlap analysis results for the common downregulated gene sets by MsigDB. **D.** GSEA NES plot for Hallmark_G2M_Checkpoint for RBBP4 g1 and g2. **E.** Heatmap of the common genes found in top three pathways based on LFC values from RNAseq results. **F**. RT-qPCR for downregulated genes in top three downregulated pathways. P values were determined by Two-way Anova compared to NT1; *p < 0.05, **p < 0.01, ***p < 0.001.

### RBBP4 knock-out results in cell cycle deformity

To explore how RBBP4 loss affects the cell cycle, we performed cell cycle analysis in U87MG-TR and U373-TR cells at PT12. We observed significant accumulation of cells in G2/M phase with RBBP4 gRNAs compared to NT1 (**Fig. 7A, Supp. Fig. 7A**), corroborating with our RNA sequencing results. Next, we investigated the effect of RBBP4 KO on the morphology of U87MG-TR cells through immunofluorescence staining (**Fig. 7B**). Staining with an antibody targeting endogenous RBBP4, we observed that RBBP4 mainly localized in the nucleus. We observed significant decrease in nuclear RBBP4 signal in KO cells compared to NT1 (**Fig. 7B, C**). The cell size increased significantly when RBBP4 was knocked out (**Fig. 7B, D**). Together with the aberrant cell size, RBBP4 KO cells exhibited nuclear defects and multinucleation, likely due to mitotic slippage events (**Supp. Fig. 8A**, **Fig. 7E**). To assess the effects of RBBP4 KO on cell growth over time, we labelled RBBP4 KO and NT1 U87MG-TR cells with H2B-mCherry and performed live cell imaging for 48 hours, capturing images every 30 minutes. Representative images from 0 and 48 hours demonstrated the reduced confluency and the number of mCherry positive cells in KO cells compared to NT1, indicating a slower growth rate upon RBBP4 KO (**Fig. 7F**). To further explore the growth rate differences, we examined the duration of mitosis. While NT1 cells spent approximately 50 minutes in mitosis, this period increased to over 100 minutes in RBBP4 KO cells, indicating a significant delay (**Fig. 7G**). Formation of multinucleated cell upon failed mitosis was also evident in live cell imaging (**Supp. Fig. 8B**). To conclude, our findings indicate that RBBP4 knock-out leads to G2/M arrest followed by increased cell size, longer mitosis and multinucleation, highlighting its critical role in cell cycle progression in TMZ-resistant cells.

**Figure 7:**
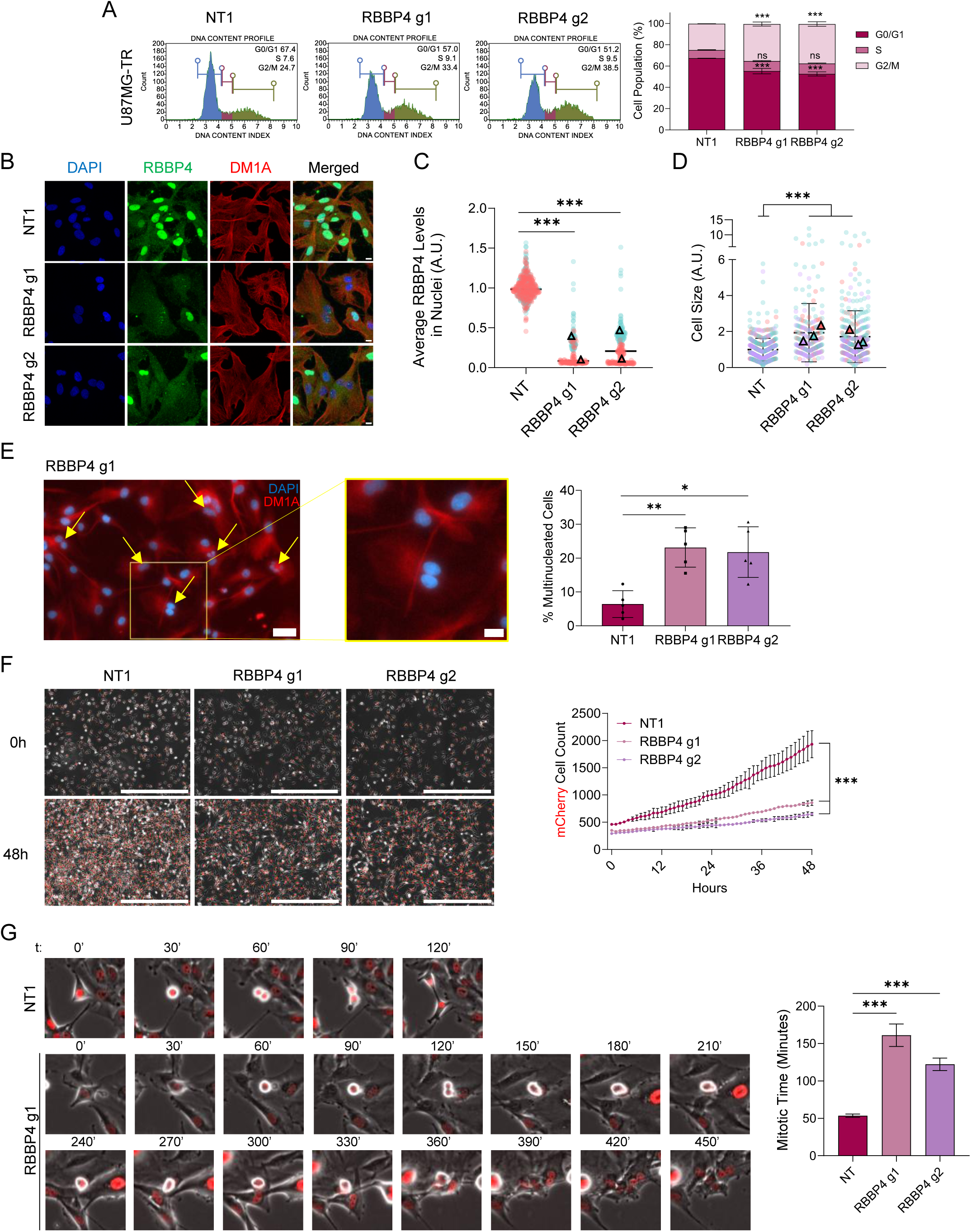
RBBP4 KO results in increased cell size and nuclear defects in U87MG-TR cells. **A.** Cell cycle analysis of RBBP4 knock-out at PT12. P values were determined by Two-way Anova between NT1-gRNA; *p < 0.05, **p < 0.01, ***p < 0.001. **B.** Immunofluorescence staining of RBBP4 (green) and DM1A (red) of U87MG-TR cells upon RBBP4 KO (PT9) (Scale bar = 20 µM). **C**. RBBP4 immunofluorescence intensity. Quantification was performed using CellProfiler. P values were determined by Mann-Whitney test between NT1-gRNA; *p < 0.05, **p < 0.01, ***p < 0.001. **D**. Cell size analysis upon RBBP4 KO. Quantification was performed using CellProfiler. P values were determined by Mann-Whitney test between NT1-gRNA; *p < 0.05, **p < 0.01, ***p < 0.001. **E**. Representative image of multinucleation, arrows indicate the multinucleated cells in the frame. (Scale bar = 50 and 20 µM). Quantification of multinucleation by CellProfiler, P values were determined by Mann-Whitney test between NT1-gRNA; *p < 0.05, **p < 0.01, ***p < 0.001. **F**. Representative images of 48h live cell with H2B-mCherry labelled U87MG-R cells upon RBBP4 KO (Scale bar = 1000 µM). P values were determined by Two-way Anova between NT1-gRNA at each data point; *p < 0.05, **p < 0.01, ***p < 0.001. **G.** Mitotic time analysis of NT1 and gRNAs. Images show the time frames where mitosis begins, and cytokinesis occurs. Quantification was performed with ImageJ. P values were determined by Mann-Whitney test between NT1-gRNA; *p < 0.05, **p < 0.01, ***p < 0.001.

## DISCUSSION

In this study, we aimed to identify chromatin modifiers critical for therapy resistance and cell survival in glioblastoma. For this purpose, we generated two TMZ-resistant glioblastoma cell lines from U87MG and U373 cells, utilizing common resistance generation mechanisms as dose escalation (U87MG-TR) and pulse treatment (U373-TR) methods (40). Both models exhibited key resistance traits, such as increased IC50 values, sustained colony-forming ability, and unchanged cell cycle dynamics under TMZ pressure. U87MG-TR cells formed tumors *in vivo* that were unresponsive to TMZ and displayed high proliferative capacity, as indicated by Ki67 staining, compared to naïve U87MG tumors. Transcriptome analysis highlighted the upregulation of pathways involved in cell proliferation, motility, adhesion (41) and neurogenesis, supporting the observed tumor morphology. MGMT, a known regulator of TMZ resistance, was significantly upregulated and knockout of MGMT restored TMZ sensitivity, confirming its role in resistance in both models of acquired TMZ resistance.

To explore epigenetic mechanisms underlying TMZ resistance, we conducted an epigenome-wide CRISPR/Cas9 screen using EPIKOL. We identified PCGF6, SET, MGMT, and APEX1 as potential sensitizers based on TMZ vs Control comparisons, commonly depleted in U87MG-TR and U373-TR cell lines. APEX1 is associated with DNA damage repair (42) and TMZ response in glioblastoma (43). PCGF6 is involved in stem cell regulation (44,45) and SET regulates various signaling pathways (46) and molecular processes, including microsatellite instability (47). While MGMT knockout resulted in significant TMZ sensitization, underscoring its central role in resistance mechanisms, PCGF6, SET and APEX1 did not produce expected effects. These discrepancies may result from various factors, such as off-target effects that compromise the specificity of results or inefficient gene knockouts that fail to produce a detectable phenotype (48,49). Despite these challenges, the efficacy of our positive (RPL9) and negative controls (NT1), and successful validation of MGMT as a proof-of-concept gene confirmed the reliability of the screen. These findings suggest that improving gene selection strategies may enhance the identification of actionable targets.

Next, we further investigated genes critical for cell survival in TMZ-resistant glioblastoma models in the absence of TMZ. By focusing on cancer-specific genes and excluding hits from the normal human astrocyte (NHA) screen, we identified seven common epigenetic vulnerabilities across U87MG-TR and U373-TR models: TOP2A, PCNA, RBBP4, BAP1, NSL1, EXOCS2, and CDK2. CDK2 is a serine/threonine kinase that is involved in cell cycle regulation (50), DNA damage and replication (51), and implicated for irradiation response in glioblastoma (52). TOP2A, a critical component of the transcription machinery (53), has been associated with glioblastoma invasion (54). While the role of BAP1 in glioblastoma is unclear, BAP1 functions as a ubiquitin C-terminal hydrolase (UCH), acting as a tumor suppressor alongside BRCA1 and is implicated in various cancers (55,56). EXOSC2, a subunit of the RNA exosome, has been linked to several diseases (57), including, more recently, breast cancer (58). NSL1 is a member of the MIS12 kinetochore complex involved in cell division (59) with its tumor suppressor role studied in Drosophila (60). PCNA, a pivotal player in DNA replication (61) and a known marker for cell proliferation, is considered a potential target in various cancers (62). Among these, Retinoblastoma Binding Protein 4 (RBBP4) emerged as a compelling candidate due to its known role as an important histone chaperone (39) and its involvement in various chromatin remodeling complexes linking it to TMZ sensitivity (26–28). Knockout of RBBP4 led to decreased colony formation, impaired cell growth, and increased apoptosis in both resistant models. GFP competition assay confirmed the long-term effect of RBBP4 loss leading to decreased cell viability. Interestingly, despite the 92% homology with RBBP4 (39), RBBP7 knockout showed minimal impact on cell growth, indicating a unique role for RBBP4 in glioblastoma.

Previous studies suggest that RBBP4 influences therapy resistance through either the p300-RBBP4-MGMT axis (26,27) or an MGMT-independent pathway involving the MRN complex (28). To explore this, we knocked out RBBP4 in a parental (U87MG), an acquired TMZ-resistant (U87MG-TR) and an innately TMZ-resistant (T98G) cell line. In parental U87MG cells, which lack MGMT expression, RBBP4 KO reduced colony-forming ability in control conditions and further sensitized the cells to TMZ. Remarkably, in TMZ-resistant U87MG-TR and T98G cells, RBBP4 KO completely abolished colony formation even in the absence of TMZ. We found decreased protein and mRNA levels of MGMT upon RBBP4 KO in T98G cells, but not in U87MG-TR, suggesting that MGMT expression is not the main link for RBBP4’s role in sustaining the viability of TMZ-resistant cells. Similarly, no significant changes in MRN complex mRNA levels were observed in either resistant model upon RBBP4 loss. Altogether, these findings suggest that RBBP4 regulates the survival of TMZ-resistant cells through a distinct mechanism, independent from MGMT and alternative to those previously reported in the literature.

Transcriptome analysis of RBBP4 knockout in U87MG-TR cells revealed downregulation of G2/M checkpoint, E2F targets, and mitotic spindle pathways, implicating cell cycle dysregulation. RBBP4 is one of the five members of multi-vulval class B (MuvB) complex along with LIN9, LIN37, LIN52 and LIN54, and interaction with different proteins/complexes determines its role in cell cycle (63,64). RBBP4-containing MuvB complex regulates cell cycle progression at G2/M phase (65). Conversely, its interaction with DREAM (dimerization partner, RB-like, E2F and multi-vulval class B (MuvB)) complex mediates transcriptional repression at G0/G1 (66). RBBP4’s involvement in DREAM and MuvB complexes indicates its vital role in cell cycle progression. Notably, several downregulated genes identified in our transcriptome analysis upon RBBP4 KO, including CENPF, AURKA and KIF2C, are directly regulated by MuvB complex in G2/M phase (67). Indeed, knockout of RBBP4 led to G2/M arrest which resulted in an increase in cell size and time spent in mitosis. Failed or prolonged divisions resulted in multinucleation, demonstrating the essential role of RBBP4 in cell cycle machinery.

In conclusion, we successfully generated TMZ-resistant cell lines that exhibited resistance both *in vitro* and *in vivo*, with MGMT upregulation playing a key role in the acquisition and maintenance of resistance. Using the EPIKOL screen, we uncovered RBBP4 as a key epigenetic regulator of cell survival in resistant cells. RBBP4’s regulation of TMZ resistance was mediated through changes in cell cycle pathways, independent of its known DNA repair interactions. Collectively, these findings highlight RBBP4 as a novel epigenetic vulnerability in acquired TMZ-resistance, positioning it as a promising therapeutic target to overcome resistance in recurrent glioblastoma.

## Supporting information

Supplementary Informations

Supplementary Video for U87MG-TR NT1 Live Cell Imaging

Supplementary Video for U87MG-TR RBBP4 g1 Live Cell Imaging

Supplementary Video for U87MG-TR RBBP4 g2 Live Cell Imaging

## Author Contributions

Study design: TB-O, EYK; data generation: EYK, FS, EE, AC, EY, IK, AHDK, IK; data analysis: ACA, ADC, EY, HS, MP, APC; data interpretation: EYK, TB-O, OY-B; initial manuscript draft: EYK, TB-O, OY-B; approved final manuscript: all authors.

## Conflict of Interest

A.P.C and M.P are co-founders of Caeruleus Genomics Ltd and are inventors on several patents related to sequencing technologies filed by Oxford University Innovations. All other authors declare no conflict of interest.

## Acknowledgments

Financial support was obtained from The Scientific and Technological Research Council of Turkey (TUBITAK) (1003-216S461 Grant). We thank Defne Yigci and Eylul Sevinc Sevinc for their technical support. Representative figures were generated with BioRender.com and licensed for publication. (Fig1: TF285PVTO2, Fig3: MZ285PW5KA, Fig4: RW285Q0SBD, Supp. Fig1: GX285PW0YH, Supp. Fig3: CP285PW9DM, Supp. Fig4: YK285PWGPV, Supp. Fig5: JD285PWL7G.) The authors gratefully acknowledge the use of the services and facilities of the Koç University Research Center for Translational Medicine (KUTTAM). A.P.C. is a recipient of a Medical Research Council (MRC) career development fellowship (MR/V010182/1).

## REFERENCES

1. Louis DN, Perry A, Wesseling P, Brat DJ, Cree IA, Figarella-Branger D, et al. The 2021 WHO classification of tumors of the central nervous system: A summary. Neuro Oncol. 2021 Aug 1;23(8):1231–51.

2. Price M, Ballard C, Benedetti J, Neff C, Cioffi G, Waite KA, et al. CBTRUS Statistical Report: Primary Brain and Other Central Nervous System Tumors Diagnosed in the United States in 2017–2021. Neuro Oncol [Internet]. 2024 Oct 6;26(Supplement_6):vi1–85. Available from: https://academic.oup.com/neuro-oncology/article/26/Supplement_6/vi1/7797290

3. Wesolowski JR, Rajdev P, Mukherji SK. Temozolomide (Temodar). American Journal of Neuroradiology. 2010;31(8):1383–4.

4. Wen PY, Weller M, Lee EQ, Alexander BM, Barnholtz-Sloan JS, Barthel FP, et al. Glioblastoma in adults: A Society for Neuro-Oncology (SNO) and European Society of Neuro-Oncology (EANO) consensus review on current management and future directions. Vol. 22, Neuro-Oncology. Oxford University Press; 2020. p. 1073–113.

5. Lee SY. Temozolomide resistance in glioblastoma multiforme. Genes Dis [Internet]. 2016;3(3):198–210. Available from: 10.1016/j.gendis.2016.04.007

6. Tomar MS, Kumar A, Srivastava C, Shrivastava A. Elucidating the mechanisms of Temozolomide resistance in gliomas and the strategies to overcome the resistance. Vol. 1876, Biochimica et Biophysica Acta - Reviews on Cancer. Elsevier B.V.; 2021.

7. Woo PeterYM, Li Y, Chan AnnaHY, Ng StephanieCP, Loong HerbertHF, Chan DannyTM, et al. A multifaceted review of temozolomide resistance mechanisms in glioblastoma beyond O-6-methylguanine-DNA methyltransferase. Glioma. 2019;2(2):68.

8. Singh N, Miner A, Hennis L, Mittal S. Mechanisms of temozolomide resistance in glioblastoma - a comprehensive review. Vol. 4, Cancer Drug Resistance. OAE Publishing Inc.; 2021. p. 17–43.

9. Villano JL, Seery TE, Bressler LR. Temozolomide in malignant gliomas: Current use and future targets. Cancer Chemother Pharmacol. 2009;64(4):647–55.

10. Mansouri A, Hachem LD, Mansouri S, Nassiri F, Laperriere NJ, Xia D, et al. MGMT promoter methylation status testing to guide therapy for glioblastoma: Refining the approach based on emerging evidence and current challenges. Neuro Oncol. 2019 Feb 14;21(2):167–78.

11. Hegi ME, Diserens AC, Gorlia T, Hamou MF, De Tribolet N, Weller M, et al. MGMT gene silencing and benefit from temozolomide in glioblastoma. New England Journal of Medicine. 2005;352(10):997–1003.

12. Ortiz R, Perazzoli G, Cabeza L, Jiménez-Luna C, Luque R, Prados J, et al. Temozolomide: An Updated Overview of Resistance Mechanisms, Nanotechnology Advances and Clinical Applications. Curr Neuropharmacol. 2020 Jun 26;19(4):513–37.

13. Gusyatiner O, Hegi ME. Glioma epigenetics: From subclassification to novel treatment options. Semin Cancer Biol. 2018;51(November):50–8.

14. Yu W, Zhang L, Wei Q, Shao A. O6-Methylguanine-DNA Methyltransferase (MGMT): Challenges and New Opportunities in Glioma Chemotherapy. Vol. 9, Frontiers in Oncology. Frontiers Media S.A.; 2020.

15. Liu L, Gerson SL. Targeted modulation of MGMT: Clinical implications. Vol. 12, Clinical Cancer Research. 2006. p. 328–31.

16. Romani M, Pistillo MP, Banelli B. Epigenetic targeting of glioblastoma. Front Oncol. 2018;8(OCT):1–9.

17. Liu A, Hou C, Chen H, Zong X, Zong P. Genetics and epigenetics of glioblastoma: Applications and Overall Incidence of IDH1 Mutation. Front Oncol. 2016;6(JAN):1–9.

18. Marampon F, Megiorni F, Camero S, Crescioli C, McDowell HP, Sferra R, et al. HDAC4 and HDAC6 sustain DNA double strand break repair and stem-like phenotype by promoting radioresistance in glioblastoma cells. Cancer Lett. 2017 Jul 1;397:1–11.

19. Kunadis E, Lakiotaki E, Korkolopoulou P, Piperi C. Targeting post-translational histone modifying enzymes in glioblastoma. Vol. 220, Pharmacology and Therapeutics. Elsevier Inc.; 2021.

20. Kayabolen A, Yilmaz E, Sahin GN, Seker-Polat F, Cingoz A, Isik B, et al. Orthogonal targeting of KDM6A/B and HDACs mediates potent therapeutic effects in IDH1-mutant glioma. Available from: 10.1101/2020.11.26.400234

21. Banelli B, Carra E, Barbieri F, Würth R, Parodi F, Pattarozzi A, et al. The histone demethylase KDM5A is a key factor for the resistance to temozolomide in glioblastoma. Cell Cycle. 2015;14(21):3418–29.

22. Banelli B, Daga A, Forlani A, Allemanni G, Marubbi D, Pistillo MP, et al. Small molecules targeting histone demethylase genes (KDMs) inhibit growth of temozolomide-resistant glioblastoma cells. Oncotarget. 2017;8(21):34896–910.

23. Yedier-Bayram O, Gokbayrak B, Kayabolen A, Aksu AC, Cavga AD, Cingöz A, et al. EPIKOL, a chromatin-focused CRISPR/Cas9-based screening platform, to identify cancer-specific epigenetic vulnerabilities. Cell Death Dis. 2022;13(8).

24. Ozyerli-Goknar E, Kala EY, Aksu AC, Bulut I, Cingöz A, Nizamuddin S, et al. Epigenetic-focused CRISPR/Cas9 screen identifies (absent, small, or homeotic)2-like protein (ASH2L) as a regulator of glioblastoma cell survival. Cell Communication and Signaling. 2023 Dec 1;21(1).

25. Zhan Y, Yin A, Su X, Tang N, Zhang Z, Chen Y, et al. Interpreting the molecular mechanisms of RBBP4/7 and their roles in human diseases (Review). Int J Mol Med. 2024 May 1;53(5).

26. Kitange GJ, Mladek AC, Schroeder MA, Pokorny JC, Carlson BL, Zhang Y, et al. Retinoblastoma Binding Protein 4 Modulates Temozolomide Sensitivity in Glioblastoma by Regulating DNA Repair Proteins. Cell Rep [Internet]. 2016;14(11):2587–98. Available from: 10.1016/j.celrep.2016.02.045

27. Mladek AC, Yan H, Tian S, Decker PA, Burgenske DM, Bakken K, et al. RBBP4-p300 axis modulates expression of genes essential for cell survival and is a potential target for therapy in glioblastoma. Neuro Oncol [Internet]. 2022 Aug 1;24(8):1261–72. Available from: https://academic.oup.com/neuro-oncology/article/24/8/1261/6540482

28. Li J, Song C, Gu J, Li C, Zang W, Shi L, et al. RBBP4 regulates the expression of the Mre11-Rad50-NBS1 (MRN) complex and promotes DNA double-strand break repair to mediate glioblastoma chemoradiotherapy resistance. Cancer Lett [Internet]. 2023;557(January):216078. Available from: 10.1016/j.canlet.2023.216078

29. Bagga M, Kaur A, Westermarck J, Abankwa D. ColonyArea : An ImageJ Plugin to Automatically Quantify Colony Formation in Clonogenic Assays. 2014;9(3):14–7.

30. Dobin A, Davis CA, Schlesinger F, Drenkow J, Zaleski C, Jha S, et al. STAR: ultrafast universal RNA-seq aligner. Bioinformatics [Internet]. 2013 Jan 1;29(1):15–21. Available from: 10.1093/bioinformatics/bts635

31. Patro R, Duggal G, Love MI, Irizarry RA, Kingsford C. Salmon provides fast and bias-aware quantification of transcript expression. Nat Methods [Internet]. 2017;14(4):417–9. Available from: 10.1038/nmeth.4197

32. Soneson C, Love MI, Robinson MD. Differential analyses for RNA-seq: transcript-level estimates improve gene-level inferences [version 2; peer review: 2 approved]. F1000Res. 2016;4(1521).

33. Love MI, Huber W, Anders S. Moderated estimation of fold change and dispersion for RNA-seq data with DESeq2. Genome Biol [Internet]. 2014;15(12):550. Available from: 10.1186/s13059-014-0550-8

34. Subramanian A, Kuehn H, Gould J, Tamayo P, Mesirov JP. GSEA-P: a desktop application for Gene Set Enrichment Analysis. Bioinformatics [Internet]. 2007 Dec 1;23(23):3251–3. Available from: 10.1093/bioinformatics/btm369

35. Onder TT, Kara N, Cherry A, Sinha AU, Zhu N, Bernt KM, et al. Chromatin-modifying enzymes as modulators of reprogramming. Nature [Internet]. 2012;483(7391):598–602. Available from: 10.1038/nature10953

36. Li W, Xu H, Xiao T, Cong L, Love MI, Zhang F, et al. MAGeCK enables robust identification of essential genes from genome-scale CRISPR/Cas9 knockout screens. Genome Biol. 2014;15(12):554.

37. Senbabaoglu F, Aksu AC, Cingoz A, Seker-Polat F, Borklu-Yucel E, Solaroglu İ, et al. Drug Repositioning Screen on a New Primary Cell Line Identifies Potent Therapeutics for Glioblastoma. Front Neurosci. 2020;14(December):1–13.

38. Leica Biosystems. H&E Staining Overview: A Guide to Best Practices [Internet]. [cited 2023 Dec 12]. Available from: https://www.leicabiosystems.com/knowledge-pathway/he-staining-overview-a-guide-to-best-practices/

39. Cai L, Liu B, Cao Y, Sun T, Li Y. Unveiling the molecular structure and role of RBBP4/7: implications for epigenetic regulation and cancer research. Front Mol Biosci. 2023 Nov 13;10.

40. McDermott M, Eustace AJ, Busschots S, Breen L, Crown J, Clynes M, et al. In vitro development of chemotherapy and targeted therapy drug-resistant cancer cell lines: A practical guide with case studies. Front Oncol. 2014;4 MAR.

41. González-Mariscal L, Miranda J, Gallego-Gutiérrez H, Cano-Cortina M, Amaya E. Relationship between apical junction proteins, gene expression and cancer. Biochimica et Biophysica Acta (BBA) - Biomembranes [Internet]. 2020;1862(9):183278. Available from: https://www.sciencedirect.com/science/article/pii/S0005273620301097

42. Fung H, Demple B. A Vital Role for Ape1/Ref1 Protein in Repairing Spontaneous DNA Damage in Human Cells. Mol Cell [Internet]. 2005 Feb 4;17(3):463–70. Available from: 10.1016/j.molcel.2004.12.029

43. Ströbel T, Madlener S, Tuna S, Vose S, Lagerweij T, Wurdinger T, et al. Ape1 guides DNA repair pathway choice that is associated with drug tolerance in glioblastoma. Sci Rep [Internet]. 2017;7(1):9674. Available from: 10.1038/s41598-017-10013-w

44. Yang CS, Chang KY, Dang J, Rana TM. Polycomb Group Protein Pcgf6 Acts as a Master Regulator to Maintain Embryonic Stem Cell Identity. Sci Rep [Internet]. 2016;6(1):26899. Available from: 10.1038/srep26899

45. Lan X, Ding S, Zhang T, Yi Y, Li C, Jin W, et al. PCGF6 controls neuroectoderm specification of human pluripotent stem cells by activating SOX2 expression. Nat Commun [Internet]. 2022;13(1):4601. Available from: 10.1038/s41467-022-32295-z

46. Westermarck J, Hahn WC. Multiple pathways regulated by the tumor suppressor PP2A in transformation. Trends Mol Med [Internet]. 2008;14(4):152–60. Available from: https://www.sciencedirect.com/science/article/pii/S1471491408000452

47. Wang H, Qiu P, Zhu S, Zhang M, Li Y, Zhang M, et al. SET nuclear proto-oncogene gene expression is associated with microsatellite instability in human colorectal cancer identified by co-expression analysis. Digestive and Liver Disease [Internet]. 2020;52(3):339–46. Available from: https://www.sciencedirect.com/science/article/pii/S1590865819307224

48. Bock C, Datlinger P, Chardon F, Coelho MA, Dong MB, Lawson KA, et al. High-content CRISPR screening. Vol. 2, Nature Reviews Methods Primers. Springer Nature; 2022.

49. Doench JG. Am i ready for CRISPR? A user’s guide to genetic screens. Nat Rev Genet [Internet]. 2018;19(2):67–80. Available from: 10.1038/nrg.2017.97

50. Fagundes R, Teixeira LK. Cyclin E/CDK2: DNA Replication, Replication Stress and Genomic Instability. Vol. 9, Frontiers in Cell and Developmental Biology. Frontiers Media S.A.; 2021.

51. Tadesse S, Anshabo AT, Portman N, Lim E, Tilley W, Caldon CE, et al. Targeting CDK2 in cancer: challenges and opportunities for therapy. Drug Discov Today [Internet]. 2020;25(2):406–13. Available from: https://www.sciencedirect.com/science/article/pii/S135964461930460X

52. Wang J, Yang T, Xu G, Liu H, Ren C, Xie W, et al. Cyclin-Dependent Kinase 2 Promotes Tumor Proliferation and Induces Radio Resistance in Glioblastoma. Transl Oncol [Internet]. 2016;9(6):548–56. Available from: https://www.sciencedirect.com/science/article/pii/S1936523316301073

53. Roca J. Topoisomerase II: a fitted mechanism for the chromatin landscape. Nucleic Acids Res [Internet]. 2009 Feb 1;37(3):721–30. Available from: 10.1093/nar/gkn994

54. Hu Y, Xue Z, Qiu C, Feng Z, Qi Q, Wang J, et al. Knockdown of NUSAP1 inhibits cell proliferation and invasion through downregulation of TOP2A in human glioblastoma. Cell Cycle [Internet]. 2022 Sep 2;21(17):1842–55. Available from: 10.1080/15384101.2022.2074199

55. Carbone M, Harbour JW, Brugarolas J, Bononi A, Pagano I, Dey A, et al. Biological Mechanisms and Clinical Significance of BAP1 Mutations in Human Cancer. Cancer Discov [Internet]. 2020 Aug 3;10(8):1103–20. Available from: 10.1158/2159-8290.CD-19-1220

56. Caporali S, Butera A, Amelio I. BAP1 in cancer: epigenetic stability and genome integrity. Discover Oncology [Internet]. 2022;13(1):117. Available from: 10.1007/s12672-022-00579-x

57. Morton DJ, Kuiper EG, Jones SK, Leung SW, Corbett AH, Fasken MB. The RNA exosome and RNA exosome-linked disease. RNA [Internet]. 2018 Feb 1;24(2):127–42. Available from: http://rnajournal.cshlp.org/content/24/2/127.abstract

58. Lv CG, Cheng Y, Zhang L, Wu GG, Liang CY, Tao Z, et al. EXOSC2 Mediates the Pro-tumor Role of WTAP in Breast Cancer Cells via Activating the Wnt/β-Catenin Signal. Mol Biotechnol [Internet]. 2024;66(9):2569–82. Available from: 10.1007/s12033-023-00834-8

59. Kline SL, Cheeseman IM, Hori T, Fukagawa T, Desai A. The human Mis12 complex is required for kinetochore assembly and proper chromosome segregation. Journal of Cell Biology [Internet]. 2006 Apr 3;173(1):9–17. Available from: 10.1083/jcb.200509158

60. Clemente-Ruiz M, Muzzopappa M, Milán M. Tumor suppressor roles of CENP-E and Nsl1 in Drosophila epithelial tissues. Cell Cycle [Internet]. 2014 May 1;13(9):1450–5. Available from: 10.4161/cc.28417

61. Boehm EM, Gildenberg MS, Washington MT. Chapter Seven - The Many Roles of PCNA in Eukaryotic DNA Replication. In: Kaguni LS, Oliveira MT, editors. The Enzymes [Internet]. Academic Press; 2016. p. 231–54. Available from: https://www.sciencedirect.com/science/article/pii/S1874604716300051

62. Wang SC. PCNA: a silent housekeeper or a potential therapeutic target? Trends Pharmacol Sci [Internet]. 2014;35(4):178–86. Available from: https://www.sciencedirect.com/science/article/pii/S0165614714000303

63. Litovchick L, Sadasivam S, Florens L, Zhu X, Swanson SK, Velmurugan S, et al. Evolutionarily Conserved Multisubunit RBL2/p130 and E2F4 Protein Complex Represses Human Cell Cycle-Dependent Genes in Quiescence. Mol Cell [Internet]. 2007 May 25;26(4):539–51. Available from: 10.1016/j.molcel.2007.04.015

64. Fischer M, Müller GA. Cell cycle transcription control: DREAM/MuvB and RB-E2F complexes. Vol. 52, Critical Reviews in Biochemistry and Molecular Biology. Taylor and Francis Ltd; 2017. p. 638–62.

65. Sadasivam S, Duan S, DeCaprio JA. The MuvB complex sequentially recruits B-Myb and FoxM1 to promote mitotic gene expression. Genes Dev [Internet]. 2012 Mar 1;26(5):474–89. Available from: http://genesdev.cshlp.org/content/26/5/474.abstract

66. Sadasivam S, Decaprio JA. The DREAM complex: master coordinator of cell cycle-dependent gene expression [Internet]. Vol. 13, NATURE REVIEWS | CANCER. 2013. Available from: www.nature.com/reviews/cancer

67. Engeland K. Cell cycle arrest through indirect transcriptional repression by p53: I have a DREAM. Vol. 25, Cell Death and Differentiation. Nature Publishing Group; 2018. p. 114–32.

