## Supplementary Informations for "Functional Genomic Screens Reveal RBBP4 as a Key Regulator of Cell Cycle Progression in TMZ-Resistant Glioblastoma"

### Supplementary Information

#### Supplementary Tables

| Supplementary Table 1: Antibodies | | |
|---|---|---|
| **Name** | **Brand** | **Cat. #** |
| MGMT | Cell Signaling Technologies | 2739S |
| GAPDH | Elabscience | E-AB-40516 |
| RBBP4 (RBAP48) | Proteitech | 20364-1-AP |
| Alpha-Tubulin | Abcam | ab15246 |
| Alpha-Tubulin | Elabscience | E-AB-20069 |
| Goat anti-rabbit | Abcam | ab97051 |

| Supplementary Table 2: qRT-PCR Primers | | |
|---|---|---|
| **Name** | **Forwar Primer** | **Reverse Primer** |
| GAPDH | AGCCACATCGCTCAGACAC | GCCCAATACGACCAAATCC |
| MGMT | GTG ATT TCT TAC CAG CAA TTA GCA | CTG CTG CAG ACC ACT CTG TG |
| RBBP4 | ATGACCCATGCTCTGGAGTG | GGACAAGTCGATGAATGCTGAAA |
| RAD50 | GGAAGAGCAGTTGTCCAGTTACG | GAGTAAACTGCTGTGGCTCCAG |
| MRE11 | ATGCAGTCAGAGGAAATGATACG | CAGGCCGATCACCCATACAAT |
| NBN | GACTGGCGTTGAGTACGTTGT | TGATTTCGGCTGATCGACTGA |
| TOP2A | GTGGCAAGGATTCTGCTAGTCC | ACCATTCAGGCTCAACACGCTG |
| PLK1 | GCACAGTGTCAATGCCTCCAAG | GCCGTACTTGTCCGAATAGTCC |
| CENPF | AGCACGACTCCAGCTACAAGGT | CATCATGCTTTGGTGTTCTTTCTG |
| AURKA | GCAACCAGTGTACCTCATCCTG | AAGTCTTCCAAAGCCACTGCC |
| MKI67 | TGTGCCTGCTCGACCCTACA | TGAAATAGCGATGTGACATGTGCT |
| DLGAP5 | AGCTGCTAATGAAAACGAACCAGAA | TCCATCTGGATTCATTCCACTTGTT |
| PIF1 | CGAGCCTAGCACAGAAGCC | CCCAGGATTCGCTTTAGCAG |
| CDKN3 | TCCGGGGCAATACAGACCAT | GCAGCTAATTTGTCCCGAAACTC |
| CCNB2 | CAACCCACCAAAACAACA | AGAGCAAGGCATCAGAAA |

| | | |
|---|---|---|
| NEK2 | TTG GAG CAG AAA GAA CAG GAG C | TCC CCA CTG AAA TGA ACT TTC TTC |
| CDK2 | ATGGATGCCTCTGCTCTCACTG | CCCGATGAGAATGGCAGAAAGC |
| SET | AGCAAGAAGCGATTGAACACA | TGGTTGGCGGAGTTTGTTATATT |
| NSL1 | CCGCTTCGTGCAAAAGCTC | TCCAGGATCTTTCTGGGATACTG |
| BAP1 | GCTCGTGGAAGATTTCGGTGT | TCATCAATCACGGACGTATCATC |
| EXOSC2 | CAATCACTACGGACACAGGATTC | TTCGTCCCACTACGATGTCTC |
| PCNA | ACACTAAGGGCCGAAGATAACG | ACAGCATCTCCAATATGGCTGA |
| APEX1 | CAATACTGGTCAGCTCCTTCG | TGCCGTAAGAAACTTTGAGTGG |
| RPL9 | GCACAGTTATCGTGAAGGGC | TTACCCCACCATTTGTCAACC |
| PCGF6 | TGAAGGCACGGGACATTTTAAG | TATTGCACGTCGGATTTCCCT |
| RBBP7 | TCTGCGGATAAGACCGTAGC | GGCGGTCAGTACCACTTGAA |

| **Supplementary Table 3: gRNA Sequences** | | |
|---|---|---|
| **Epikol Rank** | **Name** | **Sequence** |
| 3675 | MGMT-g1-F | CACCGGGTACTTGGAAAAATGGACA |
| | MGMT-g1-R | AAACTGTCCATTTTTCCAAGTACCC |
| 3680 | MGMT-g2-F | CACCGTGAAATGAAACGCACCACAC |
| | MGMT-g2-R | AAACGTGTGGTGCGTTTCATTTCAC |
| 5109 | RBBP4_g1_F | CACCGTCTTGGAACCCAAATCTCAG |
| | RBBP4_g1_R | AAACCTGAGATTTGGGTTCCAAGAC |
| 5110 | RBBP4_g2_F | CACCGTCTTTGTTGCGATGATACAA |
| | RBBP4_g2_R | AAACTTGTATCATCGCAACAAAGAC |
| 216 | APEX1-g1-F | CACCGCCAAGAGCAGATCTTGAGTG |
| | APEX1-g1-R | AAACCACTCAAGATCTGCTCTTGGC |
| 2064 | EXOSC2-g2-F | CACCGCGACACTAAGAAACATCTAG |
| | EXOSC2-g2-R | AAACCTAGATGTTTCTTAGTGTCGC |
| 628 | BAP1-g1-F | CACCGTCAAATGGATCGAAGAGCGC |
| | BAP1-g1-R | AAACGCGCTCTTCGATCCATTTGAC |
| 7439 | PCNA-2_F | CACCGTGCTTCAAATACTAGCGCCA |

| | | |
|---|---|---|
| | PCNA-2_R | AAACTGGCGCTAGTATTTGAAGCAC |
| 1104 | CDK2-g1-F | CACCGCAAATATTATTCCACAGCTG |
| | CDK2-g1-R | AAACCAGCTGTGGAATAATATTTGC |
| 4174 | NSL1-1_F | CACCGCCGCTCTGCGAGATGCGCAG |
| | NSL1-1_R | AAACCTGCGCATCTCGCAGAGCGGC |
| 6533 | TOP2A_g1_F | CACCGATTCAGTACCAAATTTACTG |
| | TOP2A_g1_R | AAACCAGTAAATTTGGTACTGAATC |
| 7593 | RPL9-2_F | CACCGATGACTACAAATAGTCCGAA |
| | RPL9-2_R | AAACTTCGGACTATTTGTAGTCATC |
| 5124 | RBBP7_g1_F | CACCGCAGGATTACATTCTCCACTT |
| | RBBP7_g1_R | AAACAAGTGGAGAATGTAATCCTGC |
| 5455 | SET-g2-F | CACCGCCGGCCGCACCATGTGGCGG |
| | SET-g2-R | AAACCCGCCACATGGTGCGGCCGGC |
| 4377 | PCGF6-g2-F | CACCGGGGAGGCCGGCAGGACTCGG |
| | PCGF6-g2-R | AAACCCGAGTCCTGCCGGCCTCCCC |

### Supplementary Figures

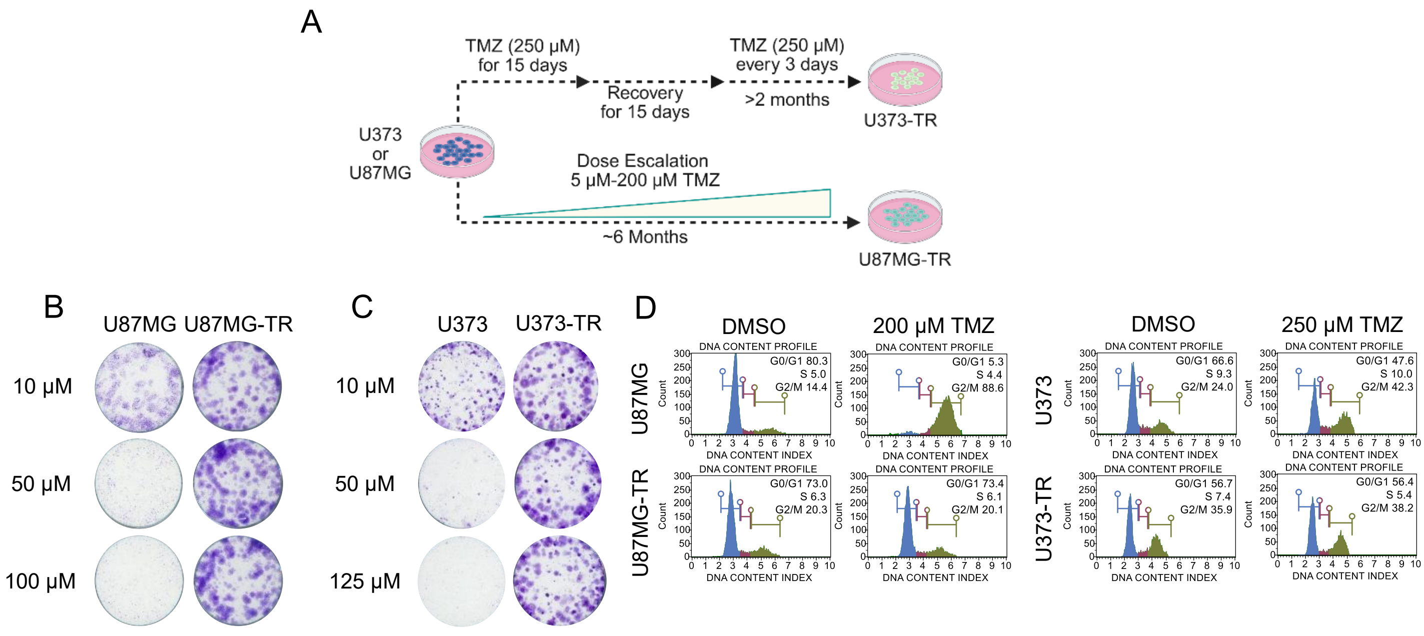


**Supplementary Figure 1: Establishment of TMZ-resistance.** **A.** Schematics of generation of resistant cell lines. Dose escalation method was performed for U87MG starting from 5 µm up to 200 µm, and high dose (250 µm) pulse method was used for U373 cell lines. Figure generated with BioRender.com **B, C.** Long term colony formation results for increasing dosages of TMZ for naive and resistant cell lines. **D.** Histograms of cell cycle analysis for TMZ response of naive and resistant cell lines (200 µM for U87MG and U87MG-TR, 250 µm for U373 and U373-TR).

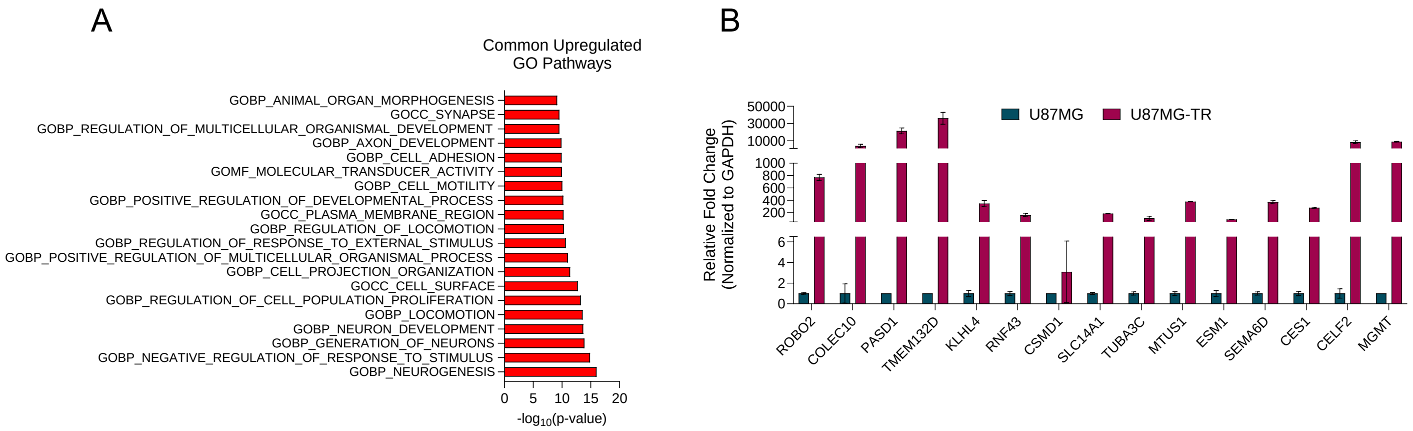


**Supplementary Figure 2**: **RNAseq analysis of parental-resistant cell lines. A.** Gene Ontology pathway analysis for 182 common upregulated genes in U87MG-TR and U373-TR. **B.** RT-qPCR analysis for the top upregulated common genes in U87MG and U87MG-TR.

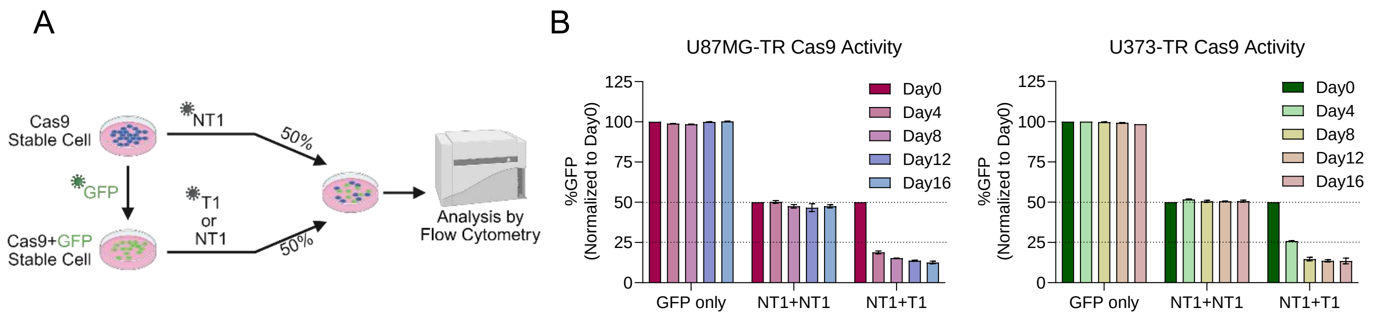


**Supplementary Figure 3: Cas9 activity of TMZ-resistant cell lines. A.** Schematics for Cas9 activity assay. Cas9 stable cells were infected with NT1 lentivirus and Cas9+GFP stable cells were infected with either NT1 or T1 lentiviruses. Following puromycin selection, cells were mixed 50%-50% and Flow Cytometry was used to measure GFP levels every four days. (Day0=Posttransfection Day5) Figure generated with BioRender.com **B.** %GFP results for U87MG-TR and U373-TR cell lines, normalized to Day0.

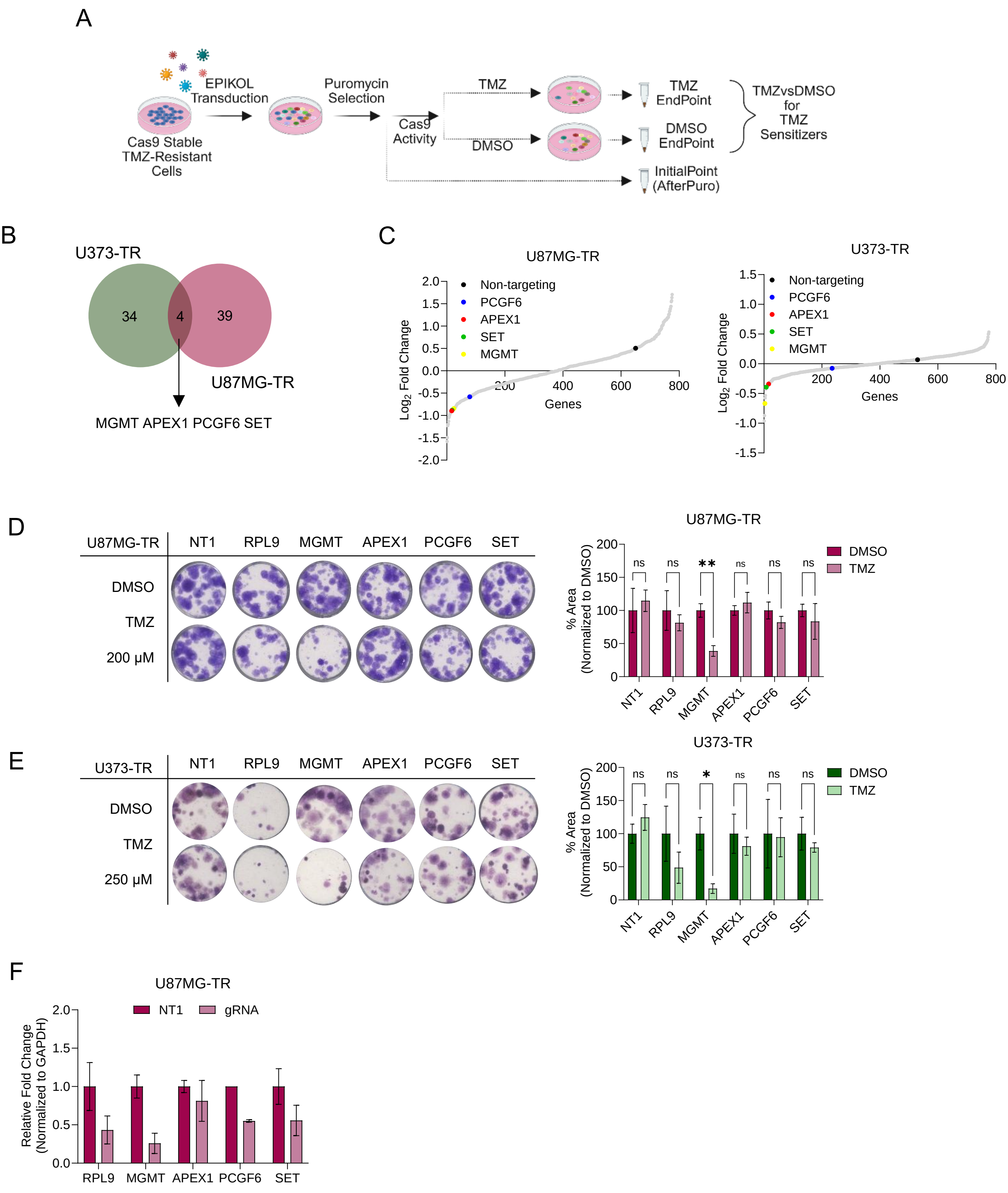


**Supplementary Figure 4: Identification of potential TMZ sensitizers based on endpoint comparisons. A**. Schematic for EPIKOL screen strategy. Cells were infected with EPIKOL lentiviruses as 1000X coverage with MOI:0.4 and selected with puromycin. Initial pellet was obtained after puromycin selection (AfterPuro). Following Cas9 activity, cells were divided into two groups as DMSO and TMZ.

Screen continued until each group reached 14-16 population doublings and endpoint pellets were collected. Endpoints were compared to identify potential TMZ sensitizers. Figure generated with BioRender.com **B.** Venn diagram for the common genes for DMSOvsTMZ comparison in two cell lines **C.** Distribution of depleted genes from DMSOvsTMZ comparison based on LFC values, selected hits are highlighted for both cell lines. **D, E.** Colony formation assay results for selected hits. Quantification of colonies were performed by ImageJ program with ColonyArea plugin, normalized to DMSO of each gRNA. P values were determined by t-test in comparison to DMSO of group; $*p < 0.05$, $**p < 0.01$, $***p < 0.001$. **F.** RT-qPCR results demonstrating downregulation of the genes targeted by gRNAs.

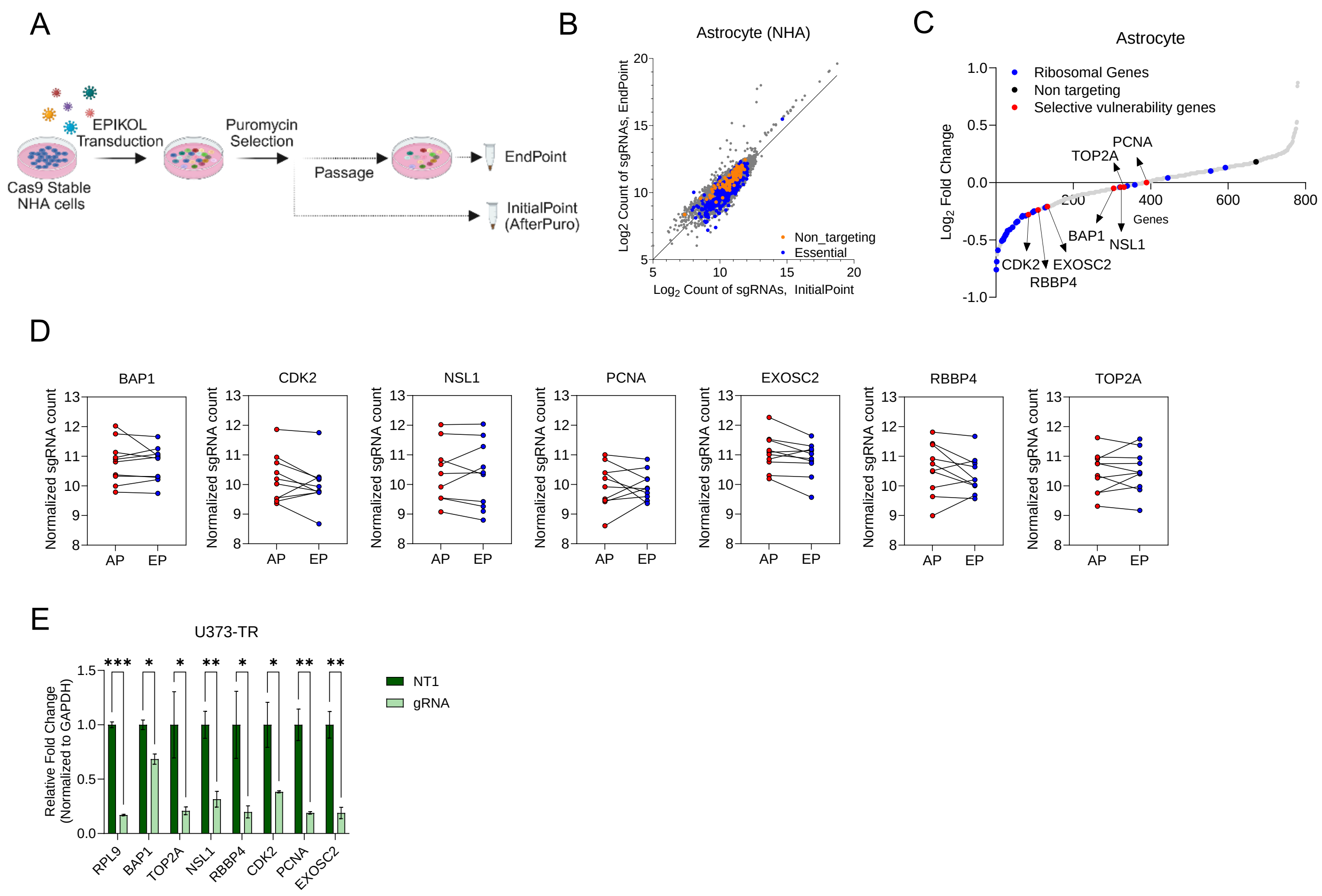


**Supplementary Figure 5: EPIKOL screen analysis and validation. A**. Schematic for EPIKOL screen strategy for NHA cells. Cells were infected with EPIKOL lentiviruses as 125X coverage with MOI:0.4 and selected with puromycin. Initial pellet was obtained after puromycin selection (AfterPuro). Screen continued for 6 weeks post transduction and endpoint pellets were collected. Figure generated with BioRender.com **B.** gRNA distribution of EPIKOL screen results for NHA. **C.** LFC graph of Astrocyte EPIKOL screen results with selected hits indicated. **D.** sgRNA count comparisons between InitialPoint (AfterPuro (AP)) and EndPoint (EP) for vulnerability genes in NHA cell line. **E.** RT-qPCR results showing downregulation of selected hits targeted with individual gRNAs. P values were determined by t-test compared to NT1; *p < 0.05, **p < 0.01, ***p < 0.001.

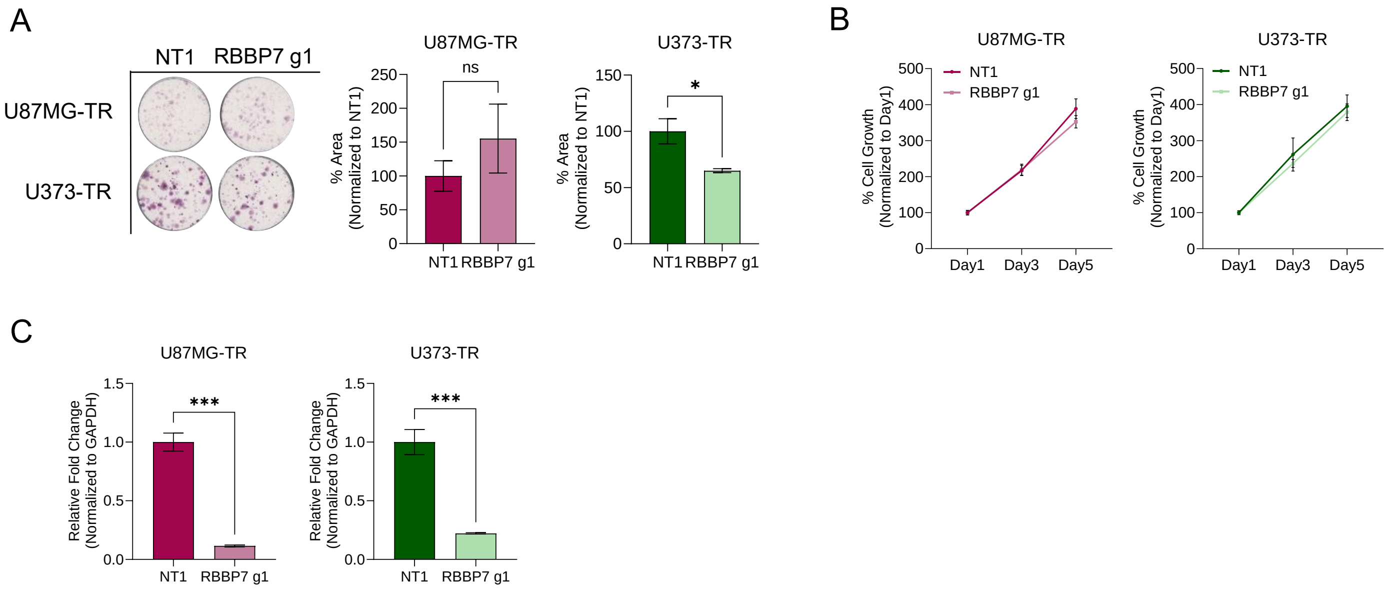


**Supplementary Figure 6: Effect of RBBP7 KO on TMZ-resistant cells. A.** Long term colony formation assay of RBBP7 knock-out and quantification of colony formation assay. P values were determined by t-test compared to NT1; *p < 0.05, **p < 0.01, ***p < 0.001. **B.** Cell proliferation curve at days 1, 3 and 5 following RBBP7 knock-out (PT10, 12, 14 respectively). **C.** RT-qPCR results showing downregulation of RBBP7 mRNA level upon KO. P values were determined by t-test compared to NT1; *p < 0.05, **p < 0.01, ***p < 0.001.

A

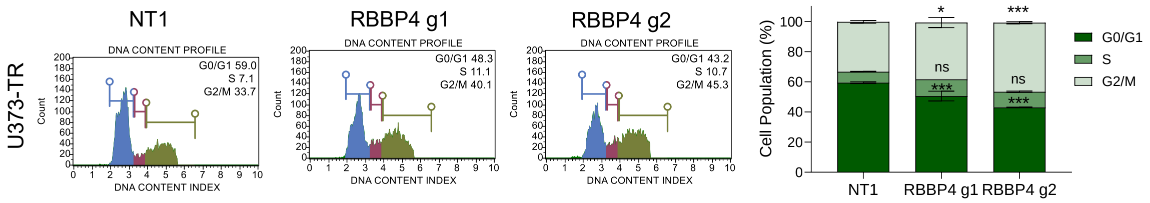


**Supplementary Figure 7: Effect of RBBP4 loss on cell cycle. A.** Cell cycle analysis of RBBP4 knock-out with U373-TR cells at PT12. P values were determined by Two-way Anova between NT1-gRNA; *p < 0.05, **p < 0.01, ***p < 0.001.

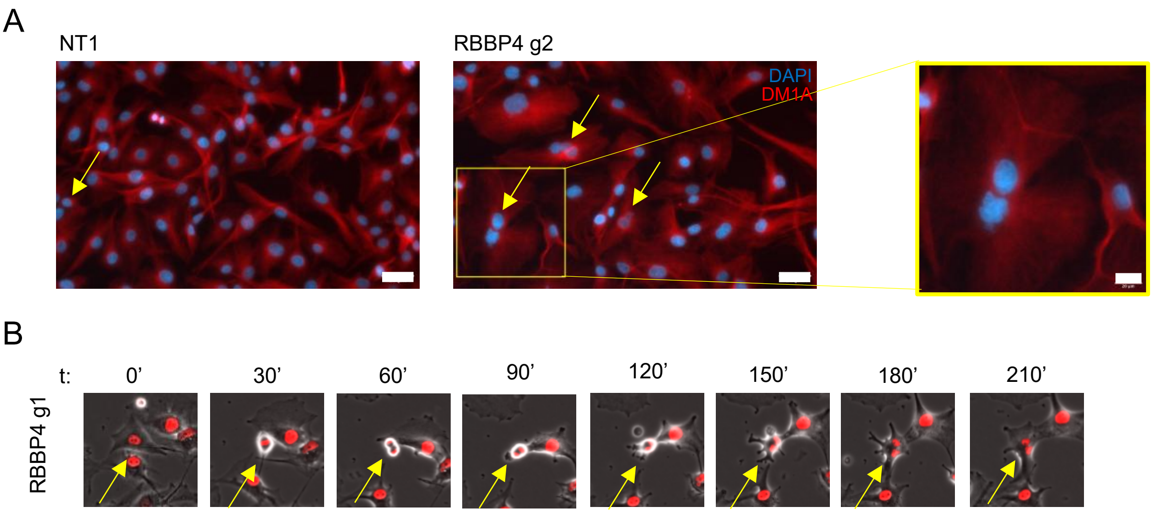


**Supplementary Figure 8: Multinucleation analysis of U87MG-TR cells upon RBBP4 KO. A.** Representative image of multinucleation, arrows indicate the multinucleated cells in the frame. (Scale bar = 50 and 20 μM). **B.** Time-framed multinucleation of a dividing cell upon RBBP4 KO.
